# Structural variants, hemizygosity and clonal propagation in grapevines

**DOI:** 10.1101/508119

**Authors:** Yongfeng Zhou, Andrea Minio, Mélanie Massonnet, Edwin Solares, Yuanda Lv, Tengiz Beridze, Dario Cantu, Brandon S. Gaut

## Abstract

Structural variants (SVs) affect plant phenotypes, but they are a largely unexplored feature of plant genomes. Little is known about the type and size of SVs, their distribution among individuals or their evolutionary dynamics. Here we identify SVs and study their evolutionary dynamics in clonally propagated grapevine cultivars and their outcrossing wild relatives. To catalog SVs, we assembled the highly heterozygous Chardonnay genome, for which one in seven genes is hemizygous. Using genomic inference as the standard, we extended SV detection to population samples. We found that negative selection acts against SVs, but particularly against inversion and translocation events. SVs nonetheless accrue as recessive heterozygotes in clonal lineages. They also define outlier regions of genomic divergence between wild and cultivated grapevines, suggesting roles in domestication. Outlier regions include the sex determination region and the berry color locus, where independent large, complex inversions drive convergent phenotypic evolution.

## MAIN TEXT

Hundreds of economically important crops are long-lived perennials. These perennials typically outcross in nature but are propagated clonally under cultivation^1^. Clonal propagation captures genotypes in a state of permanent heterozygosity that increases over time as somatic mutations accumulate^2^. To date, however, there have been few insights into heterozygous genomes, because it is technological easier to sequence either homozygous or haploid source material. The effect of this bias has been a lack of insight into the structural variants (SVs) that distinguish heterozygous chromosomes and a concomitant dearth of understanding both about the evolutionary processes that affect SVs and about their effects on phenotypes. This gap of knowledge is critical, because GWAS implicate SVs as major contributors to phenotypic variation^3,4^ and because SVs play an important role in adaptation^5^. As an example of the latter, SVs are the causative genetic variant for at least one-third of known domestication alleles^6^.

Here we study the evolutionary history and potential phenotypic effects of SVs in domesticated grapevine (*Vitis vinifera* ssp *sativa;* hereafter *‘sativa’*), a clonally propagated crop. Grapevines are arguably the most important horticultural crop in the world^7^, with ~76 million tons of fruit harvested globally in 2015^8,9^. They were domesticated from their wild ancestor, the wild Eurasian grapevine (*Vitis vinifera* ssp. *sylvestris;* hereafter *‘sylvestris’*), nearly ~8,000 years ago in the Transcaucasus^10^. Domestication increased sugar content in the berry, enlarged berry and bunch size, altered seed morphology, and prompted a shift from dioecy – i.e., separate male and female individuals – to hermaphroditism^11^. In theory, hermaphroditic grape cultivars can be selfed; in practice, selfed progeny are often non-viable. Consequently, most grape cultivars represent crosses between distantly related parents, resulting in high heterozygosity levels within cultivars^12–15^. Here our goal is to fill a major gap in our understanding of plant genome evolution by comprehensively cataloging the type and size of SVs within wild and domesticated grapevines, by inferring the evolutionary forces that shape their persistence and by investigating their potential phenotypic effects.

### SVs in Assembled Genomes

Our strategy to study the evolution of SVs was to first infer them from a phased and highly contiguous genome and then to apply the knowledge gained to a population resequencing sample. To date, however, grape genomes have been neither phased nor highly contiguous. Accordingly, we began by generating a reference genome for the Chardonnay cultivar, choosing a clone (FPS 04) that is grown worldwide. To resolve this highly heterozygous genome, we employed a hybrid sequencing approach. Hybrid assembly resulted in a contig N50 of 1.24Mb, and application of Hi-C improved the scaffold assembly N50 to 24.5Mb, vastly extending contiguity relative to other grape genomes^13,14,16,17^ (Table 1). The resulting primary assembly was 605Mb in length, which is similar to the 590Mb assembly of Cabernet Sauvignon (Cab08)^17^. The Char04 primary assembly had a BUSCO score of 93.4%, contained 38,020 annotated protein-coding genes, and consisted of 47.3% transposable elements (TEs), particularly from the *gypsy* and *copia* superfamilies (Tables 1 & **S1**).

**Table 1.** Assembly statistics of the Chardonnay genome and two comparatives: the PN40024 reference and the Cabernet Sauvignon (Cab08) assembly.

| Cultivar | Abbrev. | Assembly statistics |  |  | Annotation |  |  |
| --- | --- | --- | --- | --- | --- | --- | --- |
|  |  | Assembly size | Contig N50 (Mb) | Scaffold N50 (Mb) | #Genes | %BUSCO | %TE |
| Chardonnay | Char04<sup>1</sup> | 606 Mb | 1.24 | 24.5 | 38,020 | 93.4 | 47.3 |
| Cabernet Sauvignon | Cab08<sup>2</sup> | 591 Mb | 2.17 | - | 36,687 | 92.5 | 51.1 |
| Pinot Noir | PN40024<sup>3</sup> | 486 Mb | 0.102 | 3.4 | 41,163 | 96.9 | 47.0 |

<sup>1</sup> This paper.
<sup>2</sup> Reference<sup>17</sup>
<sup>3</sup> Reference<sup>33</sup>

We identified heterozygous SVs (hSVs) within Char04 by remapping SMRT reads to the Char04 reference^18^, revealing 18,998 hSVs of length ≥ 50 bp (Figure 1A & **Table S2**). Only 0.3% of the hSVs were detected as homozygous (**Table S2**), suggesting a low rate of misassembly. After masking these regions, observed hSVs were as long as 5.3 Mb and together constituted 91.21 Mb, or 15.1%, of the 605Mb primary assembly. hSVs were assigned to five categories relative to the reference: deletions (DELs), duplications (DUPs), inversions (INVs), translocations (TRAs), and mobile elements insertions (MEIs). DEL and MEI events were the most numerous, with 8,302 and 7,772 (**Table S2**), respectively. In addition to SVs ≥ 50 bp in length, we also detected 119,067 small (< 50bp) indels and 873,159 SNPs.

**Figure 1:**
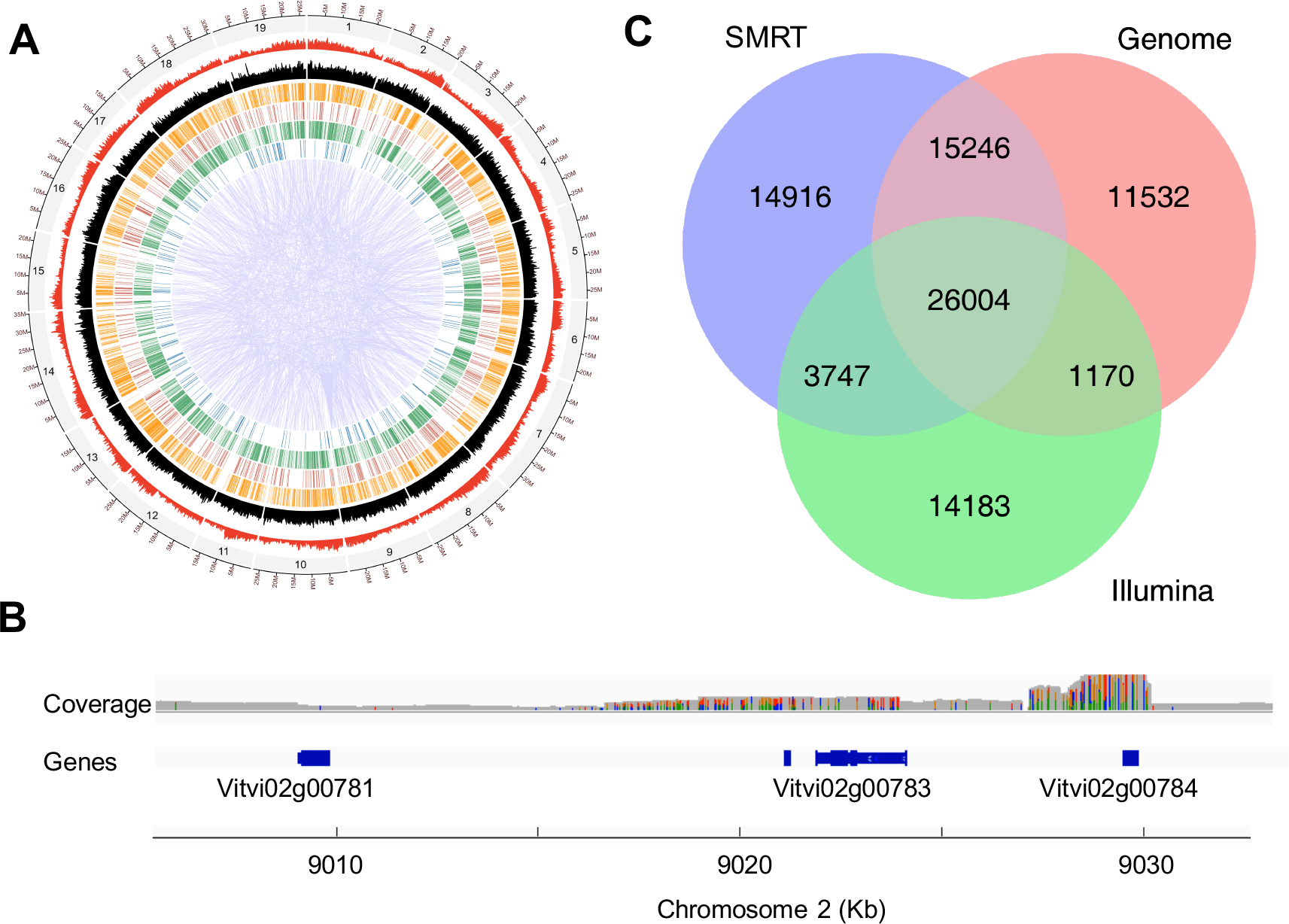
Structural heterozygosity within Chardonnay 04 and comparisons of structural variation between Chardonnay 04 and Cabernet Sauvignon 08. A) The circle plot reports heterozygous SVs within the Char04 genome. The outermost circle denotes the number and size of chromosomes (gray), followed by gene density (red), TE density (black), deletions (orange), duplications (dark red), insertions (green), inversions (blue) and with translocations represented in the middle of the circle in purple. B) A demonstration of hemizygous genes of Char04 supported by both homozygosity and coverage. The vertical colored lines in the grey coverage plot shows heterozygous sites. Both coverage and heterozygous sites support a complete hemizygous gene (Vitvi02g00781), a partial hemizygous gene (Vitvi02g00783). C) A Venn diagram showing the common and specific SVs detected by each method between Cab08 and Char04. The SVs shared between Illumina and Pacbio calls provide the basis for criteria to identify SVs from the diversity panel.

Surprisingly, 5,546 genes were hemizygous in Char04 based on inferences from long-read-mapping (Figure 1B), representing 14.6% of all annotated protein-coding genes. This value is consistent with the overall proportion of chromosomal heterozygosity by length, but it also raises concerns that it could be artificially high due to artifacts in mapping or in the Char04 reference. To allay these concerns, we performed two additional analyses to detect hSVs. First, we repeated the analysis by mapping Char04 long reads to the PN40024 reference. We detected slightly more (6,419) hemizygous genes, but they again constituted ~15% of all annotated genes in the reference. Second, we mapped SMRT reads from Cab08 to the Cab08 assembly and detected 5,702 (15.5%) hemizygous genes within this cultivar. All of these analyses are consistent in indicating that SVs affect hemizygosity for ~1 in 7 genes in cultivars.

The Cab08 assembly is less contiguous than Char04 but nonetheless permitted a rare opportunity to infer SVs based on long read sequencing between individuals from a single cultivated species^19^. We detected SVs between genomes using three approaches. We first mapped SMRT reads from Cab08 to the Char04 primary assembly (**Figure S1**). These results yielded ~3-fold higher numbers of SV events between cultivars than within Char04 (**Table S2**), reflecting the distinct parentage of Chardonnay and Cabernet Sauvignon^7,20–22^. Of 59,913 inferred SVs, DEL and MEI events were again most numerous, with 24,138 and 21,722 events, respectively, between genomes. SMRT read alignment further confirmed high hemizygosity of protein-coding genes, because the two cultivars differed in ploidy level for 9,330 genes. Of these, 2,217 exhibited presence/absence variation (PAV), similar to previous estimates based on less complete data^12,23^. Based on GO analyses, PAV genes are biased toward functions in defense response, flower development, membrane components and transcription factors (*P* < 0.001).

We also compared Char04 and Cab08 primary assemblies by whole genome alignment^24^ (**Figure S2**), which yielded a similar numbers of SVs (52,952) but fewer MEI events (**Table S2**). Finally, we mapped 25× Illumina reads from Cab08 to Char04, which detected 62% of the number of SVs based on SMRT reads (**Table S2**). The length distribution of SVs varied among the three methods; SMRT-read analyses detected more large (>10kb) events (**Figure S3**). Importantly, 75% of SVs inferred by SMRT-read alignment were confirmed by either genome alignment or short-read analyses (Figure 1C; **Figure S4**). These confirmed SVs encompassed 1,822 PAV genes and 45,403 MEIs between Char04 and Cab08, attesting to substantial SV variation among cultivars.

### Negative selection on SVs

To gain wider information about SVs in grapevines and their wild relatives, we amassed short-read sequencing data representing 50 grapevine cultivars and 19 wild relatives (**Table S3**). The application of short-read alignment for detecting SVs is subject to high levels of false-negatives and -positives^25^. To limit false-positives, we relied on our Char04 to Cab08 comparisons, specifically the subset of SVs confirmed by both long-read and short-read alignments. We examined their mapping quality, mapping depth and likelihood to provide empirical cut-offs for short-read SV calls. After applying these cut-offs to the population sample, we filtered overlapping and complex SVs to obtain a highly curated set of 481,096 SVs for population analyses (**Table S4**). These SVs yielded relationships among accessions that were remarkably similar to those based on SNPs, providing assurance about their reliability (**Figure S5**).

Given our population set of SVs, we computed the unfolded site frequency spectrum (SFS) for 12 *sylvestris* samples and a down-sampled set of 12 *sativa* samples chosen after genetic analysis (**Figures S6-S8**). The SFS for the two taxa were similar overall (Figure 2A), reflecting the fact that cultivated grapevine did not undergo a severe domestication bottleneck^7,15^ that can dramatically alters population frequencies. In both taxa, all SV types exhibited leftward shifts of the SFS relative to synonymous SNPs (sSNPs). Their SFS differed significantly from that of sSNPs in both taxa (*P* < 0.05, Kolmogorov-Smirnov, Bonferroni corrected), suggesting that SVs are predominantly deleterious.

**Figure 2:**
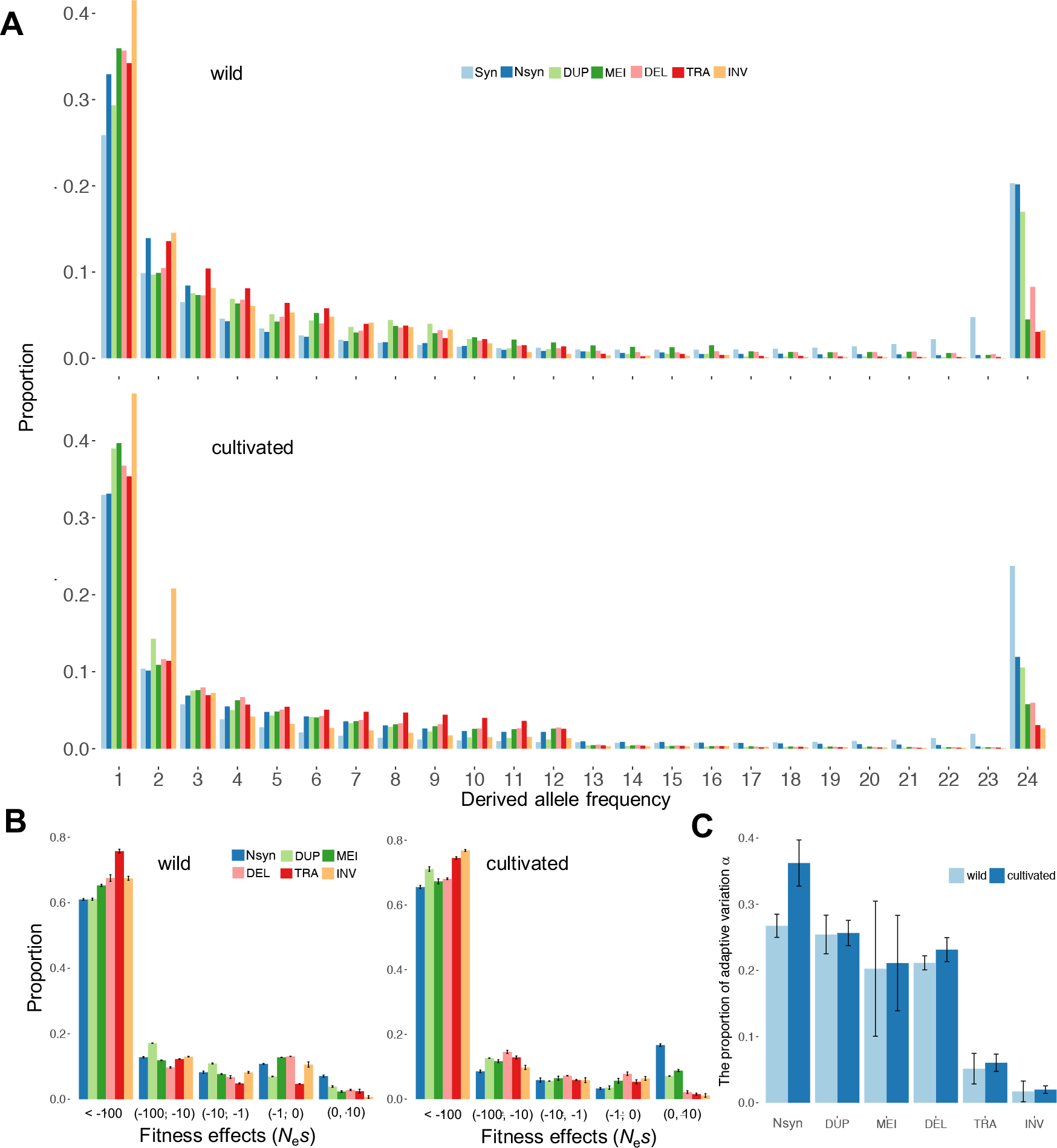
SVs are strongly deleterious and under purifying selection. A) The unfolded site frequency spectrum (SFS) of different types of SVs compared to presumably neutral synonymous SNPs (Syn) and nonsynonymous SNPs (Nsyn) for samples of 12 wild (top) and 12 cultivated (bottom) grapevines. The types of SVs plotted include duplications (DUP), TE polymorphisms (MEI), deletions (DEL), translocations (TRA) and inversions (INV). B) The inferred distribution of fitness effects (*N*_e_*s*) for SVs and nonsynonymous SNPs in wild (left) and cultivated (right) grapevines. C) The proportion of adaptive variation (*a*) in wild and cultivated grapevines.

To quantitate the strength of selection against SVs, we estimated the distribution of fitness effects (DFE) from population frequency data, using sSNPs as a neutral control. In both taxa, the results confirmed that non-synonymous SNPs (nSNPs) and SVs undergo strong purifying selection (Figure 2B). They also revealed variation among SV types, because TRA events and INV events were more strongly selected against in both taxa, mirroring their more extreme SFSs. These inferences were also consistent with estimates of α, the proportion of adaptive variants, because α was estimated to be lower for INVs (<2%) and for TRAs (<7%) than for DUP (α=25% for *sylvestris*), DEL (α=21%) and MEI (α=20%) events (Figure 2C). α estimates for SVs were lower than those based on nSNPs (27% and 36% for *sylvestris* and *sativa*, respectively), which were comparable to other perennial taxa^26^. Based on DFE and α estimates, negative selection appears to be stronger in *sativa* than *sylvestris* (Figure 2). However, the comparison between taxa must be interpreted with caution because the inferential models were designed to analyze outcrossing species like *sylvestris* and not clonally propagated crops. Nonetheless, the results strongly suggest that SV events are more deleterious than nSNPs, on average, and that INV and TRA events are especially deleterious.

### SVs accumulate in clonal propagants

SVs are deleterious, on average, but clonal propagation may allow variants to hide as heterozygous recessives^15,27^. The accumulation of recessive mutations was evident from three aspects of *sativa* genetic diversity. First, within individual heterozygosity was 11% higher, on average, within *sativa* than *sylvestris* based on SNPs (**Figure S9**). Second, sheltering of recessive mutations was evident from calculations of the additive SV load, which is the number of number of heterozygous mutations plus twice the number of derived homozygous mutations per individual^28^. Individual cultivars have a 6% higher additive SV load than their wild counterparts, on average, due to elevated heterozygosity (Figure 3A). Enhanced load was not evident for homozygous SVs or for presumably neutral sSNPs (Figure 3A), suggesting that deleterious SVs accrue and are sheltered in the heterozygous state. These patterns of SV load are consistent with forward simulations showing that clonal propagation can lead to the accumulation of deleterious recessive mutations without a notable fitness decrease^15^. Finally, the SFS provided evidence of sheltering of recessive mutations within *sativa*, based on the marked reduction in frequency for any variants over 50% (Figures 2A&**S8**). This unexpected observation may have a simple explanation: when a variant has a frequency over 50% in a clonally propagated population, then at least one individual must be homozygous, so that the recessive variant is exposed to negative selection.

**Figure 3:**
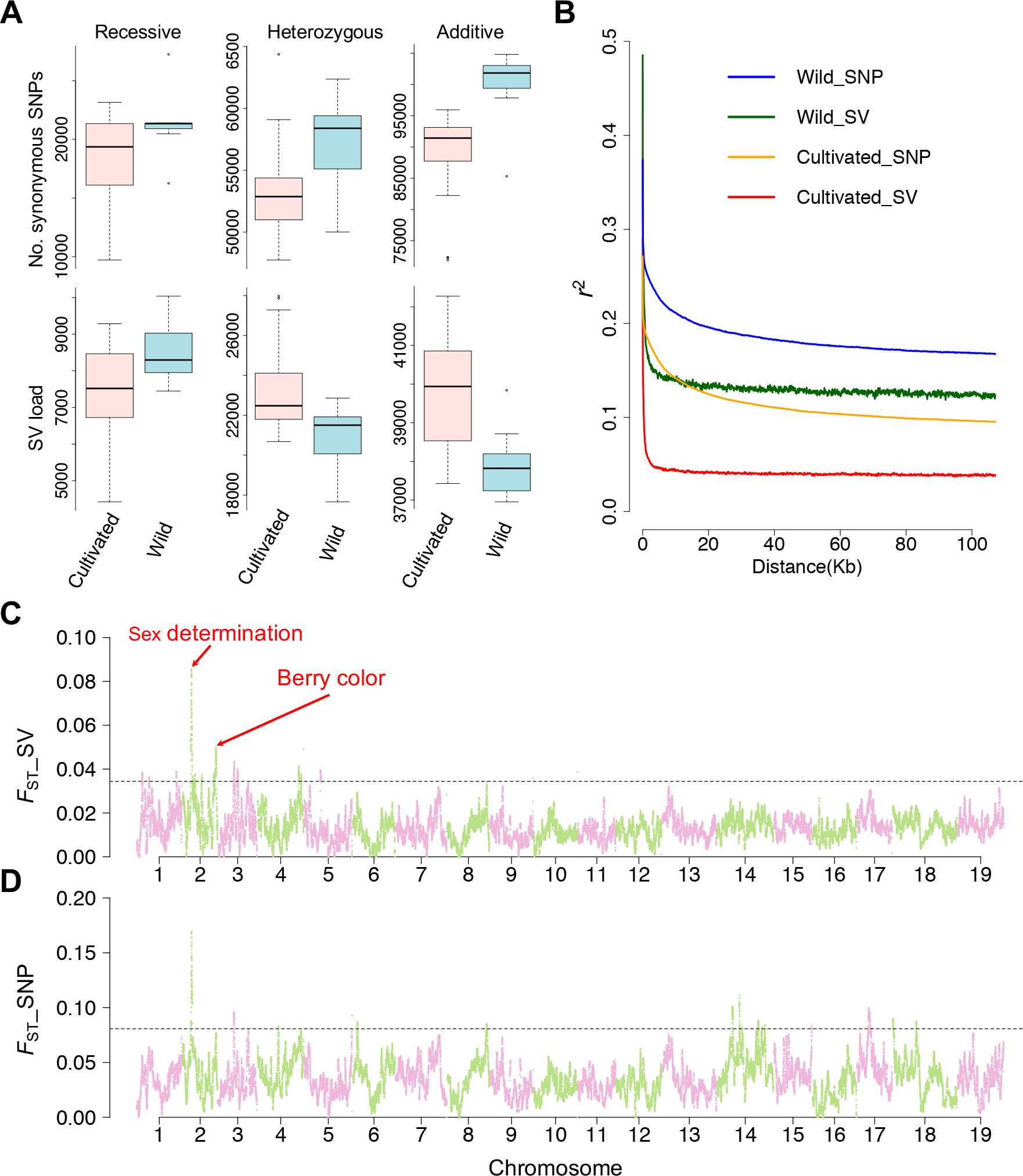
Population genetics of SVs associated with grapevine domestication. A) The recessive (number of homozygous SVs per grapevine), heterozygous and additive (the number of heterozygous SVs plus two times the number of homozygous SVs per grapevine) load in wild and cultivated grapevines for SVs compared to presumably neutral sSNPs. B) The decay of LD, as measured by *r*^2^, of SVs and SNPs as a function of physical distances between markers. C) Genetic differentiation between *sylvestris* (n = 12) and *sativa* (n = 50) sample across the genome, based on F_ST_ of SVs within 20 kb sliding windows. The dashed horizontal line represents the cut-off for the 1% tail of the F_ST_ distribution. Peaks of divergence corresponding to the sex region and the berry color loci are indicated. The *x-*axis indicates the number and size of chromosomes across the genome. D) The same as panel C, except genetic differentiation is based on SNP data.

The accumulation of heterozygous variants should affect linkage disequilibrium (LD), both because LD decreases as a function of population frequency^29^ and because cultivated grapes tend to have more low frequency variants than their wild counterparts (Figure 2A). Consistent with this observation, LD decays more rapidly over physical distance for *sativa* than for *sylvestris*, despite the relative dearth of recombination via outcrossing in cultivars. LD also decays more rapidly for SVs than for SNPs in both taxa. This last finding is important because SVs have been implicated to affect phenotypes and explain more phenotypic variation than SNPs^3,4^. However, the more rapid decline of LD for SVs suggests that it may be difficult to identify causative SVs by relying on linkage and also that reliance on SNPs for association mapping is likely to miss SVs that affect phenotypes.

### SV outliers, domestication and sex

Cultivated grapevine differs phenotypically from its wild relatives^11^. In theory, the genes that contribute to these phenotypes can be inferred from population genetic data as regions of marked chromosomal divergence between wild and cultivated samples. We estimated both SNP and SV divergence across the genome, as measured by F_ST_ in fixed windows of 20 kb (Figure 3C). Overall, average F_ST_ estimates were substantially higher for SNPs (0.0354 ± 0.0165) than SVs (0.0135 ± 0.0066), reflecting that individual SVs are typically found at lower population frequencies (Figure 2A).

We ranked the top 1% (or 485) F_ST_ windows for both SNPs and SVs. SNP-based windows generally conformed to a previous study^15^, but SNPs and SVs both identified QTL regions on chromosome 2 that correspond to the sex-determination region and to the berry color locus (Figure 3C). An additional 410 SV-based windows were found on chromosomes 1, 2, 3, 4, and 5. Of these 410, only 81 (19.8%) overlapped with windows that also had significantly higher F_ST_ for SNP divergence. Based on GO analyses, high F_ST_ windows were enriched for a few functional classes, including stilbenoid and folate biosynthesis. Stilbenes are particularly interesting because they accumulate in seeds and berry skin during berry ripening, vary in concentration between cultivars, and include resveratrol^30^, a component thought to have beneficial effects on human health. We also detected 78 diagnostic (or fixed) SVs between wild and cultivated samples that were associated with the gain and loss of seven and 10 *sativa* genes, respectively (**Table S5**). Among the 10 lost, four were NBS-LRR disease resistance genes located between 11.053 to 11.064 Mb on chromosome 9 of PN40024.

The highest F_ST_ peak for SVs corresponded to the sex determination (SD) region on chromosome 2 (Figure 3C), which also contained more SV events relative to the genomic background (*P* = 0.0067; χ^2^). Mutations in the SD region caused the shift in mating system during domestication. After confirming that the sex-linked region corresponds to 4.90Mb and 5.04Mb on PN40024^31,32^ (**Figure S10**), we resolved, for the first time, complete SD haplotypes and their underlying SVs. Chardonnay is rare among cultivars because it is a homozygote for the hermaphroditic *(H*) haplotype^32^. We compared its two *H* haplotypes to the PN40024 primary assembly^33^, which is thought to represent the female (*F*) haplotype^32^. Four genes exhibited PAV variation between *H* and *F* haplotypes. One of these, *VviAPT3*, has been proposed as a candidate SD gene^31^, because it may have a role in the abortion of pistil structures^34^. However, *VviAPT3* was present in both the *H* and *F* haplotypes of Cab08 (Figure 4A), suggesting that the lack of *VviAPT3* on PN40024 was an assembly error. The remaining three PAV genes (a *DEAD DEAH* box RNA helicase gene, the TPR-containing protein and the unknown protein previously known as *ETO1*) differentiated *H* from *F* haplotypes (Figure 4A). We also annotated two previously unrecognized genes, *Inaperturate pollen 1* (*VviINP1*) and a *C2H2*-type Zinc finger, in both *F* and *H* haplotypes. *INP1* expression in *Arabidopsis* alters the deposition of pollen apertures^35^ and could confer pollen sterility in females.

**Figure 4:**
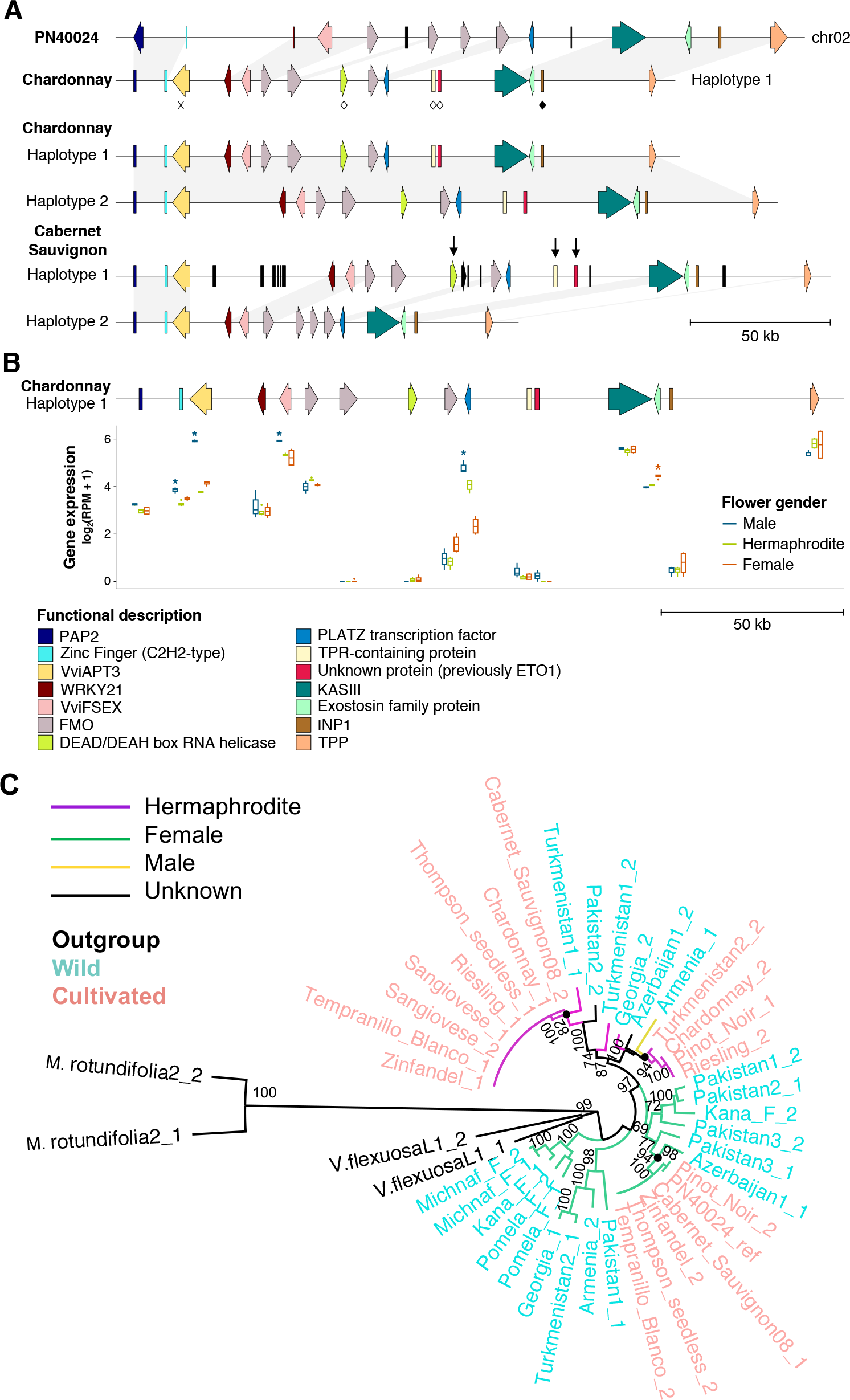
Haplotypes of the sex region and the evolution of sex in grapevine. A) Comparison of the sex determination region among cultivars. The PN40024 (V2) haplotype represents the primary assembly. Chardonnay is homozygous hermaphroditic (HH), and both haplotypes from Char04 are shown. Cabernet Sauvignon is heterozygous (HF), with Haplotype 1 of Cab08 representing the presumed H haplotype. * denotes the gene *VviAPT3* that is absent from PN40024 assembly but found in both F and H haplotypes; open diamonds denote the genes located on chromosome 0 in the PN40024 assembly, and the filled diamond denotes a novel functional annotation in Char04 (*INP1*). Protein-coding genes are colored according to their functional annotation. Genes that are not shared among genome assemblies are colored in black. Black arrows highlight genes that are found on inferred H haplotypes in Chardonnay and Cabernet Sauvignon. B) Gene expression values of each flower gender type projected on the Chardonnay protein-coding genes are shown at both G (flowers closely pressed together) and H (flowers separating, just before blooming) stages as log_2_^(RPM + 1)^. C) A phylogeny of the sex determination region recapitulates known sex types for cultivars and detects two H clades split by the single known male in the wild sample, suggesting more than one origin of the H type.

Hermaphroditism was likely to be caused by a mutation in the dominant *F* sterility gene on the male (*M*) haplotype^32,36^. The female sterility gene is unidentified, but it is likely expressed in males and knocked-down in hermaphrodites. To identify potential candidates, we performed gene expression analyses among sexes, based on expression data from two late stages of floral development (Figure 4B). The three PAV genes were lowly expressed and thus are unlikely *F* sterility candidates, but five genes differed significantly (adj. *P* ≤ 0.05) in sex-specific expression. Four were more highly expressed in males, including *VviAPT3* and the *C2H2*-type Zinc finger gene; these four constitute plausible female sterility candidates.

To investigate whether any of these candidates housed a loss-of-function SV, we built a phylogeny of the SD region, which confirmed that H haplotypes were closer to the the *M* haplotype from our single, confirmed *sylvestris* male than to F haplotypes (Figure 4C). In fact, the *M* haplotype separated two clades of *H* haplotypes, providing support for more than one origin of hermaphroditism in cultivars^32^. We estimated estimated the two clades to be 10,705 and 13,222 years old, respectively, slightly older than the accepted date of domestication. Because the *sylvestris M* haplotype was closely related to one of the Char04 haplotypes (Figure 4C), we identified SVs within and between them. Four genes were in a hemizygous state in the wild male, including the three PAV genes, and there were also three hemizygous TEs near genes, but none were obvious candidates to affect the function of the four most plausible female sterility candidates (Figure 4B). Unfortunately, the genetic mutation(s) that causes hermaphroditism and the identity of the dominant female sterility gene remain elusive, but the region underscores the dynamics of SV events and their potential relationships to a domestication phenotype.

### Convergent Inversions Contribute to Berry Color

A second region of high F_ST_ divergence between wild and cultivated grapevines encompassed the berry color region (Figure 3C). It, too, had more SVs than the genomic background (*P* = 3.3×10^−5^, χ^2^). The region is interesting because *sylvestris* has dark berries, representing the ancestral condition^11^, and because white berries originated in a subset of *sativa* cultivars. SVs have been implicated in the origin of white berries, especially a 5’*Gret1* retroelement insertion that reduces the expression of a *myb* gene (*VviMYBA1)* that regulates anthocyanin biosynthesis^37^. Subsequently, it was shown that a frameshift mutation in a second *myb* gene (*VviMYBA2)* was also necessary to cause white berries^38^. Surprisingly, these two mutations (the *Gret1* insertion and the *VviMYBA2* frameshift) are heterozygous in most grape cultivars^39^. Somatic mutations causing white grapes delete the functional *VviMYBA1* and *VviMYBA2* alleles, leaving the plant hemizygous for null alleles^40,41^.

Given the history of the *MybA* locus and the fact that it encompasses a peak of F_ST_ divergence, we investigated the region with a chromosome scale plot of Char04 reference vs. Cab08, revealing a large 4.82Mb (chr02: 12,295,113bp-17,118,777bp) inversion in Char04 (Figure 5A). This inversion was confirmed by comparison to PN40024, by the identification of discordant and split reads at the junctions (**Figure S11**), and by the lack of an inversion between Cab08 and PN40024 (Figure 5B). The Char04 inversion was bounded by *copia* elements, suggesting they played a role in its formation. The inversion encompassed the *MybA* region, but it did not affect the number of *MybA* genes because there were nine in Char04, Cab08 and PN40024. The inversion does, however, affect hemizygosity, because the entire inverted region appears to be hemizygous on the basis of read coverage and homozygosity (**Figure S11**). Thus, white berries in Chardonnay may be attributable to two related events, a large inversion on one chromosome and a simultaneous deletion on the other.

**Figure 5:**
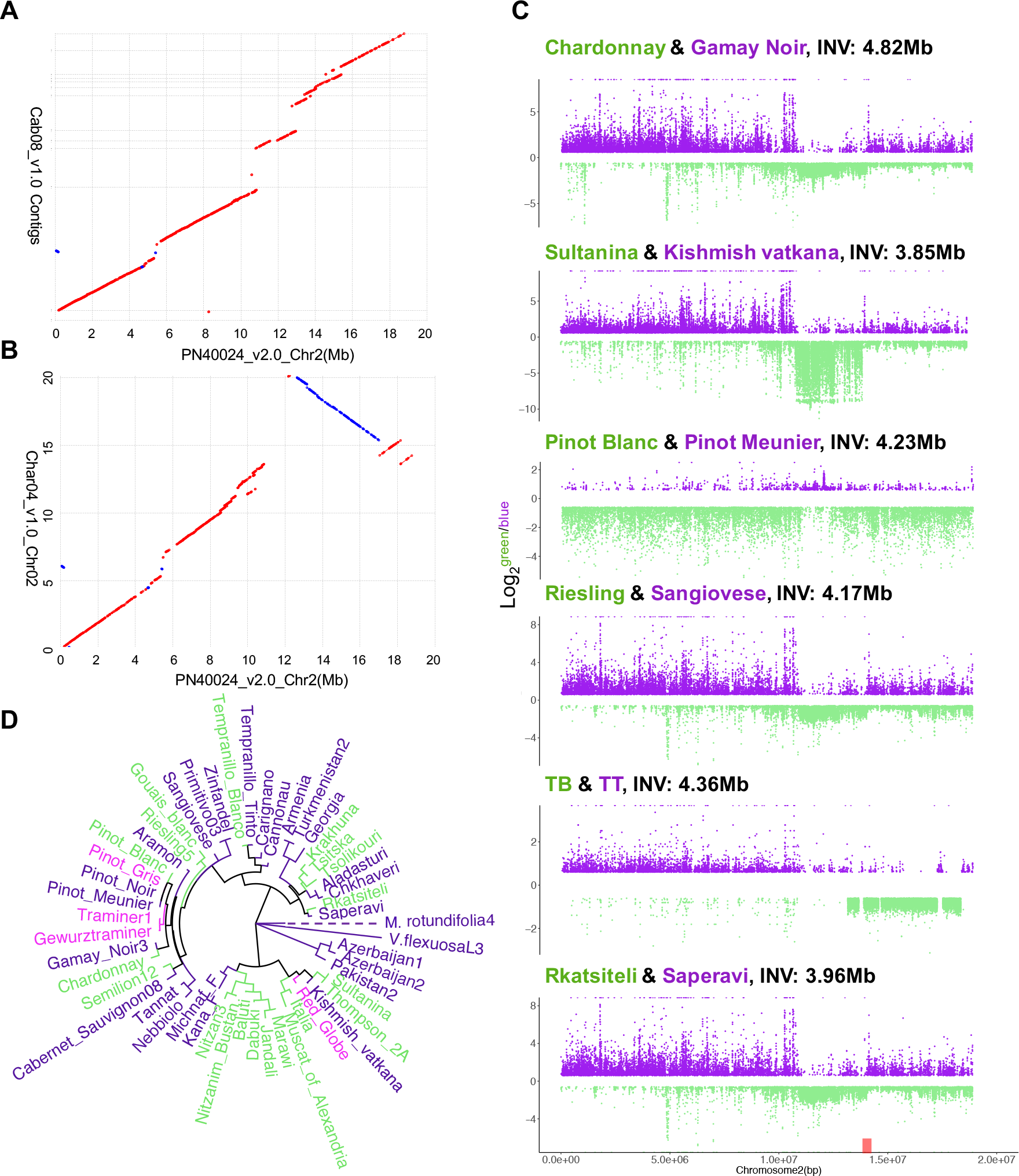
Convergent evolution of inversions associated with white berries. A) A dot plot between PN40024 chromosome 2 and Cab08 contigs. B) A dot plot between PN40024 chromosome 2 and Chard04 chromosome 2 that reveals a 4.82 Mb inversion overlapping with the major berry color QTL in grapevines. C) These plots contrast coverage across chromosome 2 for pairs of white berry and dark berry grapevines. In each contrast, the white berry grape is labeled in green. The y-axis is the log2 of white/dark read numbers so that, for example, regions of very low values indicate relatively few reads in the white-berry grape. For each contrast, the size of the inferred inversion is provided, based on the presence of split reads. TB and TT are abbreviations for Tempranillo Blanco and Tempranillo Tinto. D) A phylogeny, based on genome-wide SNPs from a selection of grape varieties, with the color of text labels reflecting the color of the berry.

Another study has recently characterized the somatic mutations that led to white berries in the Tempranillo cultivar^42^. The mutations included hemizygosity at both *VviMybA1* and *VviMybA2*, along with a series of complex series of SVs that included a putative 4.3Mb inversion on chromosome 2^42^. Given that both Chardonnay and Tempranillo have large, Mb-scale inversions associated with white berries, we investigated the generality of the association. To do so, we first built SNP-based phylogenies of white-berried cultivars and closely related dark-berried varieties. The phylogeny shows that white-berry mutations occurred independently on several occasions (Figure 5B). We then chose six pairs of closely related red and white-berried varieties and contrasted them using short-read analyses. For these short-read analyses, we focused on coverage and runs of homozygosity, while also carefully combing the data for evidence of split and discordant reads that span potential inversions. All six contrasts yielded evidence for a large inversion encompassing the *MybA* region (Figure 5C). The inferred inversions ranged from 3.85Mb to 4.82Mb in size and included from 134 to 176 genes, with 118 genes in common (including the *MybA* genes) across all six inversions. Read coverage data, which varied across pairs, strongly suggested hemizygosity of the entire inversion in at least one contrast (Sultanina vs. Kishmish vatkana) and near the *MybA* region in other contrasts (Figure 5C).

Somatic mutations to white berries are associated with hemizygosity of *MybA* genes and with large, Mb scale inversions. But why are large inversions associated with the white berry phenotype? We can think of three explanations. The first is that the inversion contains non-*MybA* genes that also affect phenotype. To investigate this hypothesis, we mapped gene expression data from red and white berries collected over four stages of berry development^43^ and counted the proportion of differentially expressed genes between color morphs. The proportion of differentially expressed genes within the Char04 inversion was no higher than the genomic background (*P* = 0.82, χ^2^), suggesting that the inversion is not enriched for genes that contribute to berry color. The second explanation is that inversions are common because of underlying properties of the chromosome 2 sequence, such as enhanced fragility^44^. The region does not contain any obvious differences in TE distribution or other gross features (Figure 1A), but this explanation remains a possibility, particularly given flanking *copia* elements in Char04. Finally, it is possible that similar inversions have occurred commonly throughout *Vitis* genome evolution, that most are lost because they are selected against (Figure 2B), but that only a few affect an obvious phenotype – like berry color – that is prone to human intervention. Whatever the underlying cause(s) for these large inversions, they represent a stunning example of convergent evolution via independent SV events.

Altogether, our sequencing of the Chardonnay genome, coupled with comparisons to the genomes of Cabernet Sauvignon and Pinot Noir, have provided insights into the evolution of clonally propagated genomes and into plant genomes more broadly. One insight is that grapevine genomes are riddled with heterozygous SVs, to the extent that they comprise up to 15% of the chromosome by length and cause 1 in 7 genes to the be hemizygous. Although negative selection acts against SVs and is particularly strong against inversions and translocations, SVs nonetheless accumulate in cultivars due to clonal propagation and the sheltering of recessive somatic mutations. Only a small proportion of these SV events are estimated to be adaptive, but some clearly associate with agronomically important phenotypes, such as hermaphroditism and white berry color. Although we cannot yet pinpoint the mutations that led to hermaphroditism, the latter originated on multiple, independent occasions via complex and large Mb-scale inversions.

## METHODS

### Genome sequencing, assembly and polishing

The Chardonnay clone chosen for sequencing was FPS 04, a clone commonly grown in California and throughout the world. The reference plant is located at Foundation Plant Services, University of California, Davis. DNA isolation and the preparation of SMRTbell libraries followed^17^. The preparation of paired-end Illumina libraries followed^15^. SMRTbell libraries were sequenced on a PacBio RSII system, generating a total of 31.51gb (52X). Illumina sequencing was conducted on a HiSeq4000 sequencing platform in 150 paired-end (PE) mode (54X) and 100 PE mode (124X). Both SMRTbell and Illumina libraries were sequenced at the UC Irvine High Throughput Genomics Center. Raw reads were deposited to the Short Read Archive (SRA) at the NCBI under the BioProject ID: PRJNAXXX.

Genome assembly was based on a hybrid strategy, that utilized both long and short sequencing reads, and that merged three separate assemblies. The first assembly utilized Canu v1.5^45^ to assemble SMRT reads, based on default parameters and with a genome size of 600 Mb. A second, hybrid assembly was generated with DBG2OLC^46^ based on contigs from the Platanus assembly and the longest 30X Pacbio reads. The Platanus assembly was based on^47^ v1.2.4 with default settings, using trimmed 178X Illumina paired-end reads. The DBG2OLC settings (options: k 31 AdaptiveTh 0.01 KmerCovTh 2 MinOverlap 30 RemoveChimera 1) were similar to those used for previous hybrid assemblies^48,49^, except that the k-mer size was increased to 31. The k-mer size was increased to minimize the number of misassemblies by including 90% of all k-mers reported by the meryl program within the Canu package^45^. The consensus stage for the DBG2OLC assembly was performed with PBDAGCON^50^ and BLASR^51^. Third, PacBio genomic reads were assembled using FALCON-Unzip v1.7.7^17^. Multiple assembly parameters (length_cutoff_pr) were tested; the least fragmented assembly was obtained with a minimum length cut-off of 9 kb. The final FALCON-Unzip parameters can be found in ***Supplemental text 1***. Unzip phasing and haplotype separation were performed with default parameters.

To integrate information obtained from the different assembly methods – Canu, DBG2OLC and FALCON-Unzip – we opted for an iterative approach of assembly merging using *quickmerge*^52^, following a broader application of assembly merging based on^49^. Quickmerge merges assemblies to increase the contiguity of the most complete (query) genome by taking advantage of the contiguity of the second reference sequence. To merge the assemblies, we followed a series of steps. First, the DBG2OLC and Canu assemblies were merged into a single assembly, QM1, using DBG2OLC assembly as the query, the Canu assembly as the reference and run options (options: hco 5.0 c 1.5 l 260000 ml 20000). Contigs that were unique to the Canu assembly were incorporated in the subsequent assembly, QM2, by a second round of *quickmerge* (options: hco 5.0 c 1.5 l 260000 ml 20000). In this second *quickmerge* run, the merged assembly from the previous step, QM1, was used as the reference assembly, and the Canu assembly was used as the query. A third round of merging (options: hco 5.0 c 1.5 l 345000 ml 20000) was performed using primary contigs of FALCON-Unzip as the reference assembly and the previous resultant assembly, QM2, as the query, generating the QM3 assembly. The final assembly, QM4, was generated by a fourth run of *quickmerge* (options: hco 5.0 c 1.5 l 345000 ml 20000), using QM3 as the reference and the Falcon-unzip assembly as the query.

All the assemblies described above, including the preliminary assemblies (Canu, DBG2OLC and Falcon-Unzip), temporary assemblies (QM1-QM3), and the final assembly (QM4), were polished twice with long reads using Quiver (Pacific Biosciences) from SMRT Analysis v2.3 (using parameter: -j 80). Long reads (> 1,000bp), consisting of ~43X coverage, were used for polishing. The assemblies were also polished twice using *Pilon* v1.16^53^ run using default settings. For this purpose, Illumina reads were aligned to the assembly using Bowtie2 v2.32^54^ and sorted using samtools v1.3^55^.

BUSCO v2.0 was used to measure gene space completeness and conserved gene model reconstruction of all generated assemblies^56^. The embryophyta database, which contained 1,440 highly conserved genes, was used to measure gene model reconstruction and estimate assembly completeness. Quast v2.3^57^ was run to calculate assembly length and N50 on each assembly. Dot plots were generated using nucmer and mumplot from MUMmer4 v3.23^24^ with the options: -l 100 -c 1000 -d 10 -banded -D 5. The BUSCO v3^56^ pipeline was applied to the final genome assembly, using the embryophyta_odb9 database.

The final assembly included both primary haplotype sequences and alternative contigs (aka haplotigs). To remove some of the alternative contigs and minimize redundancies, we performed a contig reduction. Contig reduction was executed by first aligning the final assembly to itself using Blat v. 36^58^. A python script was generated for filtering contigs that did not meet one minimum and two maximum thresholds: contig length, %query alignment and %alignment overlap. In practice, the three thresholds were investigated over ranges – e.g., minimum contig length ranged from 0, 10000, 50000, 100000 bp; % query alignment was examined over 18 randomly chosen values between 90% to 99.9999%, and % aligned overlap (PctAO) (80 and 90%), as well as maximum PctQA (100%) and PctAO (110 and 120%). New filtered genome assemblies were generated after filtering contigs based on a combinatorial of these five parameters. A gradient descent was performed on three additional parameters generated for each new filtered assembly; assembly size, contig N50 and BUSCO scores. Two formulas were generated to calculate PctQA and PctAO. 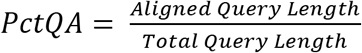 and 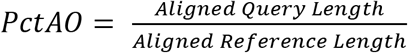. Alignments generated from contigs aligning to themselves were not considered. The scripts and code used for assembly and alternate haplotig reduction are available on GitHub: https://github.com/esolares/CAP

### Scaffolding and GapClosing

A Dovetail HiC library was prepared in a similar manner as described previously^59^. The library was sequenced on an Illumina platform to produce 211 million 2×100bp paired end reads, which provided 1,624× physical coverage of the genome (1-50kb pairs). The input *de novo* assembly, shotgun reads, and Dovetail HiC library reads were used as input data for HiRise^60^. Shotgun and Dovetail HiC library sequences were aligned to the draft input assembly using a modified SNAP read mapper (http://snap.cs.berkeley.edu). The separations of HiC read pairs mapped within draft scaffolds were analyzed by HiRise to produce a likelihood model for genomic distance between read pairs, and the model was used to identify and break putative misjoins, to score prospective joins, and make joins above a threshold. After scaffolding, shotgun sequences were used to close gaps between contigs.

MUMmer v4.0^24^ was used to identify and to sever erroneous junctions between contigs. The resulting scaffolds underwent a second scaffolding procedure using SSPACE-longreads v1.1^61^ with default parameters and a minimum coverage of 10 reads (options: -l 10). Gaps were closed using PBjelly (PBSuite v15.8.24;^62^) with default parameters for all the gap-closing steps, and assembled with options: -x ‘-w 1000000 -k – n 10’. Scaffolds were again manually curated as described above.

### Gene Annotation

Repetitive sequences were identified with RepeatMasker^63^ using the repeat library previously developed for *V. vinifera* cv. Cabernet Sauvignon^64^. *Ab initio* prediction of protein-coding genes was carried out with SNAP (ver. 2006-07-28)^65^, Augustus v3.0.3^66^, and GeneMark-ES v4.32^67^. *Ab initio* predictions were combined with the predictions of Augustus trained with BUSCO genes, as well as the gene models annotated with PASA v2.1.0^68^, using the experimental data reported in ***Supplemental text 2***. RNA-seq data obtained from public databases (***Supplemental text 2***) were *i*) assembled using both an on-genome strategy, with Stringtie v1.3.3^69^, and a *de novo* transcriptome procedure, with Trinity v2.4.0 in genome-guided mode setting a maximum intron length of 10Kb (option: --genome_guided_max_intron 10000); *ii*) clustered with CD-HIT-EST v4.6^70^, with coverage threshold 90% (option: -c 0.9); and *iii*) filtered with Transdecoder v3.0.1^71^, which retained only genes with a full-length open reading frame (ORF). Experimental evidences (transcripts and proteins) were mapped on the genome using Exonerate v2.2.0^72^and PASA v2.1.0^68^, and together with all the predictions used as input to EVidenceModeler v1.1.1^73^. Weights used in EVidenceModeler are reported in ***Supplemental text 3***. The annotation was refined and enhanced with alternative transcripts using PASA v2.1.0^73^ and assembled experimental evidences; parameters used for refining the gene structures are described in ***Supplemental text 4***. Models not showing a full-length ORF from start codon to stop codon or showing in-frame stop codons were removed. Transcripts were blast-searched for homolog proteins in the RefSeq plant protein database (ftp://ftp.ncbi.nlm.nih.gov/refseq, retrieved January 17th, 2017). Functional domains were identified using InterProScan v5^74^ using the databases provided in ***Supplemental text 5***. Gene models with no significant blast hit against RefSeq plant protein database (HSP<50 amino acids) and lacking any functional domain were discarded. Gene ontology (GO) obtained from InterPro domains and RefSeq homologs with at least 50% of reciprocal coverage and identity were combined using Blast2GO v4 (^75^ to assign a functional annotation, gene ontology (GO), and enzyme commission (EC) descriptions to each predicted transcript.

### Chromosome assignment and heterozygosity in the Chardonnay genome

The Char04 primary assembly consisted of 684 scaffolds, that summed to 606 Mb with an N50 close to that of an average grape chromosome size (25.4 Mb). We aligned the Char04 primary assembly to the PN40024 genome using the nucmer function in MUMmer4^24^. The top 23 scaffolds covered 82% (492 Mb) of the Char04 primary assembly and aligned to the PN40024 chromosomes (Fig. S1), except two long scaffolds with lengths of 20Mb (Char04v1.0_683) and 11Mb (Char04v1.0_682). These two scaffolds did not align to PN40024 genome assembly but did align to Cab08 contigs. At the same time, chromosome 13 of the PN40024 genome aligned to only a few small Char04 scaffolds. For the purposes of presentation (Figure 1).

The largest 22 scaffolds of Char04 were collinear with PN40024 and summed to 481 Mb. Each chromosome was represented by one scaffold, except chromosomes 7 and 11, which consisted of 2 and 3 scaffolds, respectively. For all ensuing analyses, we treated these 22 scaffolds as the Char04 reference genome. We evaluated heterozygosity within this reference for both small variants (SNPs + indels < 50 bp) and large structural variants (SVs ≥ 50 bp). SNPs and indels were called based on remapping 124X Illumina 100-bp PE reads to the reference. The Illumina reads for this application and for diversity analyses (see below) were trimmed using Trimmomatic-0.36 to remove adapter sequences and bases for which average quality per base dropped below 20 in 4 bp windows. Filtered reads were then mapped to the Char04 reference with default parameters implemented in bwa-0.7.12 using the BWA-MEM algorithm^76^. The bam files were filtered (unique mapping with a minimum mapping quality of 20) and sorted using samtools v1.9^55^. PCR duplicates introduced during library construction were removed with MarkDuplicates in picard-tools v1.119 (https://github.com/broadinstitute/picard). SNPs and small indels were called with the HaplotypeCaller in GATK v4.0 pipeline, and then filtered following^15^.

To identify SVs within the Char04 genome (i.e. between the two haplotypes), we called SVs using the Sniffles pipeline^18^. First, Pacbio reads longer than 500bp were mapped onto Char04 primary assembly using the two aligners Minimap2 v2.14 with the MD flag^77^ and NGMLR v0.2.7^18^, separately. Variant calling was then performed with Sniffles. SV analysis outputs (VCF files) were filtered based on the following four steps: *i*) we removed SVs that had ambiguous breakpoints (flag: IMPRECISE) and also low quality SVs that did not pass quality requirements of Sniffles (flag: UNRESOLVED); *ii*) we removed SV calls shorter than 50 bp; *iii*) we removed SVs with less than 4 supporting reads; and *iv*) we removed duplicate SV calls from Sniffles. [Sniffles frequently called multiple SVs at the same position for multiple pairs of breakpoints. In these cases, we kept the SV with the most supporting reads.] The same filtering steps were applied in downstream analyses when we called SVs between Cab08 and Char04 primary assemblies (see below). In general, using the aligner Minimap2 from the Sniffles pipeline lead to detecting more SVs (e.g., 37,169 in total within Char04) than long-read mapping with NGMLR v0.2.7 (23,972 in total within Char04). Given the differences from the two mapping protocols, we built consensus SVs calls using SURVIVOR v1.0.3^78^. Using the merged SV set, we called genotypes and combined them into a single VCF using the population calling steps of the Sniffles pipeline^18^. The genotypes of SV calls from both programs (NGMLR and Minimap2) were intersected using bedtools v2.25^79^ to get the final Pacbio SV calls. False positives associated with assembly errors were identified when homozygous no-reference (1/1) SVs were called. For downstream analyses, we masked those regions when we used Char04 primary genome assembly as the reference.

### Comparing SVs between Chardonnay and Cabernet Sauvignon

Char04 and Cab08 genomes were compared using three different alignment approaches: whole-genome alignment, long-read alignment, and short-read alignment. The first consisted then to compare primary contigs of Cab08 (N50 = 2.2 Mb) and Char04. Cab08 primary contigs were aligned to the Char04 reference using nucmer (nucmer -maxmatch -noextend) in MUMmer4^24^. After filtering 1-to-1 alignments with a minimum alignment length of 1,000 bp (delta-filter -1 -l 1000), the show-diff function and NucDiff^80^ were used to extract the features and coordinates of SVs.

The second comparison was based on alignment of SMRT reads from Cab08 onto the Char04 reference. SMRT reads from Cab08, representing ~140X coverage, were mapped onto Char04 genome using Minimap2 and NGMLR, as described above. SVs were genotyped based on merged SV calls from both mappers, using the population calling steps of Sniffles pipeline^18^. The SV calls were filtered and duplicates were removed following the four steps listed in the previous section. The genotypes of SV calls from both programs were intersected using bedtools v2.25^79^ to get the final SMRT-based SV calls. These SMRT-based SV calls were used as the “gold standard” for downstream analyses.

Finally, we mapped Cab08 Illumina PE reads corresponding to ~15X of raw coverage, which mimics the coverage of population data (see below). These reads were filtered, mapped onto the Char04 reference, and then the bam files were cleaned, sorted with PCR duplicates and masked following^15^. SVs were called with all the population samples (69 in total, see below) using both LUMPY v0.2.13^81^ and DELLY2 v0.7.7^82^. For LUMPY, the read and insert lengths were extracted from mapping files (bam files) for each sample using samtools v1.9^55^, and the SVs were genotyped using SVTyper^81^. The SV calls from DELLY and LUMPY were merged using SURVIVOR v1.0.3^78^. SVs for all 69 population samples presenting the following five criteria were retained: *i*) a minimum of three PE reads or split reads (SR) supporting the given SV event across all samples; *ii*) SV calls with precise breakpoints (flag PRECISE); *iii*) SVs passing the quality filters suggested by DELLY and LUMPY (flag PASS); *iv*) _SV length_ ≥ 50 bp; *v*) complex SVs, consisting of, or overlapping SVs were excluded. SV calls for Cab08 and Char04 were extracted using vcftools v0.1.13^83^ to permit the comparison of the three detecting methods.

The coordinates and SV features for all SV calls of Cab08 and Char04 based on whole-genome alignment, SMRT reads and Illumina short-read alignments were extracted and saved as bed files. SV calls of the three methods were compared using bedtools v2.25^79^ with a minimum reciprocal overlap of 80%. We took the intersect of the DELLY and LUMPY calls to separate SVs into three categories: *i*) shared between methods, which was roughly 74.6% of the SV calls; *ii*) DELLY-specific SVs, and *iii*) LUMPY-specific SVs. We then combined the three sets using SURVIVOR^78^ and intersected it with SMRT-based SV calls to get a shared VCF. Finally, we extracted mapping and quality statistics from the short-read SV calls that corresponded to the ‘gold standard’ long-read calls. These statistics were used in the population mappings as cut-offs to filter short-read SV calls (see below).

### SNP and SV calling for population samples

Illumina whole genome resequencing data were gathered from 69 accessions (Table S4), each of which with coverage > 10X. The mean mapping depth across accessions was 21.6X. The sample of accessions included 12 wild (ssp. *sylvestris*) samples from the Near East, where grape was domesticated, along with 50 *vinifera* cultivars that represent major lineages. The sample also included three *V. flexuosa* and four *Muscadinia rotundifolia* accessions from North America, which were used as outgroups for downstream population genetic analyses.

SNPs and indels were called for this population sample using the HaplotypeCaller in the GATK v4.0 pipeline, following^15^. SNPs and indels were filtered and annotated using SnpEff v4.0^84^, following^15^. SVs were called from short-read alignment using the LUMPY & DELLY pipelines, as described above. The merged SV genotypes were filtered following the six steps enumerated in the previous section, with the added proviso that SV calls missing in 30% of all individuals were excluded for population genetic analyses. In addition, we used statistics from the intersected set of SVs called from Cab04 to Char04 comparisons to filter ‘real’ SVs (see previous section). That is, we used statistics from the set of SVs detected by short-read alignment that were confirmed by corresponding to ‘gold standard’ SV calls by long-read alignment. These cut-off statistics included: *i*) a minimum number of supporting four reads in LUMPY calls (flag SU, which equals to SP+PE) *ii*) a minimum number of three SR or PE reads supporting each of the reference and variant alleles in DELLY calls (the flag DR/RR: number of PE/SR reads supporting the reference allele and the flag DV/RV: number of PE/SR reads supporting the variant allele); *iii*) a mapping quality ≥ 20 in DELLY calls (flag MAPQ); *iv*) a genotype quality score ≤ −5 (flag GQ) in DELLY calls. SV calls that did not pass these criteria were treated as missing data.

### Mobile element insertions (MEIs)

We used the filtered BAM files with PCR duplicates masked for each sample as input for detecting polymorphic transposable elements (TEs) with the Mobile Element Locator Tool (MELT) v2.1.4^85^. MELT uses unaligned and split reads from BWA alignments, a reference genome, and consensus TE sequences to identify polymorphic TEs. Because MELT relies on sequence similarity for identifying TEs, we used an Hidden Markov Model (HMM) method to build consensus sequences for the TE families that represented > 4% of the Char04 reference (i.e., LINES: L1; LTR retrotransposons: *Copia* and *Gypsy*; and DNA transposons: MuDR and MULE-MuDR; **Table S2**). We preprocessed BAM and TE consensus files with the Preprocess and BuildTransposonZIP utilities of MELT, respectively.

MEIs were detected across the population by using the following four steps from the MELT pipeline: *i)* TE variants compared to Char04 genome were detected for each accession individually using IndivAnalysis; *ii*) all polymorphic TE calls from all samples were merged to detect breakpoints of insertions in the reference genome using GroupAnalysis; *iii*) the resulting variants file was then used to call genotypes of all insertions for each sample using the Genotype utility; *iv*) a consensus VCF file was creating after filtering the detected MEIs using the MakeVCF utility. We again used only the first 22 longest scaffolds to represent the reference genome in these analyses, because fragmented scaffolds affect the performance of the program^85^. These four steps were performed for each TE family, separately. In order to set a threshold of maximum divergence, we used both short- and long-read alignments of Cab08 onto Char04 for calling MEI. Then, the four analysis steps were performed for each TE family, separately, with two different thresholds of maximum divergence, 5% and 10%, between putative polymorphic TEs and the consensus sequence. Comparison of the MEIs detected using short- and long-read alignments showed a higher overlap of MEIs between the two kinds of sequencing when applying a maximum divergence threshold (i.e., divergence from an inferred consensus TE) of 5% rather than 10% (58% and 33%, respectively). Accordingly, we used MEI calls based on 5% divergence for downstream analyses after filtering. MEI calls were discarded that did not pass the MELT quality filters, with imprecise breakpoints, that were missing in 30% of the population sample, and that were shorter than 50bp.

### Population genetic analyses

Our analyses of the Illumina population data resulted in SV calls for a wide variety of events, including insertions (INS), deletions (DEL), duplications (DUP), inversions (INV), and translocations (TRA). In general, variant calling using short-read alignment allowed to detect only short insertions (INS, Figure S2), and we therefore excluded INS variants from further analyses. Complex variants, which were defined as composite variant of different types (for example a reverse tandem duplicate: INVDUP), were also excluded. We also removed any DELLY & LUMPY SV calls in the remaining categories (i.e, DEL, DUP, INV, TRA) that overlapped with MEI calls or genomic regions annotated as TEs. Finally, we only retained SV calls that shared the same breakpoints across the population samples. Altogether, we considered five distinct SV categories – DEL, DUP, INV, TRA, and MEI – in our population genetic analyses. We also conducted principal component analyses (PCA) for SNP and SV calls using PLINK v1.9^86^ (Figure S6).

SNPs and SVs with a minor allele frequency > 0.1 were used for analyses of linkage disequilibrium (LD) in the wild and the cultivated grapevine samples, respectively. LD decay along physical distance were measured by the squared correlation coefficients (*r*^2^) between all pairs of SNPs within a physical distance of 300 kbp, using PLINK v1.9^86^. The decay of LD against physical distance was estimated using nonlinear regression of pairwise *r*^2^ *vs*. the physical distance between SNPs or SVs mid-positions^29^.

Since LD decayed within 20 kbp in both the wild and the cultivated samples, we divided the Char04 genome into 24,056 non-overlapping windows of 20 kbp in size to calculate genomic differentiation of SVs between wild and cultivated samples and to compare SV differentiation to SNPs. For a window to be included in downstream analyses, we required at least 1,000 bases after filtering. Levels of genetic differentiation between species at each site were estimated using the method-of-moments *F*_*ST*_ estimators based on vcftools v0.1.13^83^, which calculates indices of the expected genetic variance between and within species allele frequencies. We then averaged *F*_ST_ values of all sites within each 20 kbp non-overlapping window.

We calculated the unfolded site frequency spectrum (SFS) using the *V. flexuosa* and *Muscadinia rotundifolia* samples as outgroup. To derive the SFS, we counted the number of sites at which *k* of *n* haplotypes carry the derived variant for SNPs (synonymous: 4-fold sites, and non-synonymous sites: 0-fold sites), and SVs (DEL, DUP, INV, TRA, and MEI). To exclude direct effects of selection on synonymous sites, we detected selective sweeps based on the composite likelihood ratio (CLR) test implemented in SweeD v3.2.1^87^. Synonymous sites at genomic windows with top 5% CLR values were excluded in SFS and downstream analysis.

We calculated the SFS for the sample of 12 putatively wild *sylvestris* samples, a down-sampled set of 12 cultivars, and the full set of 50 cultivars (**Figure S7**). To identify a set of 12 cultivars to down sample, we inferred population structure across samples for all wild *sylvestris* and grapevine cultivars using the NGSadmix utility of ANGSD v0.912^88^ based on SNP sites with < 20% missing data, a minimal base quality of 20 and a minimal mapping quality of 30. We predefined the number of genetic clusters K from 2 to 8, and the maximum iteration of the Expectation Maximization (EM) algorithm was set to 10,000. Based on these population structure results (**Figure S5**), the down-sampled set of 12 cultivars was chosen to represent major genetic clusters and also to represent accessions with the least missing data (**Table S4**).

### Distribution of fitness effects (DFE) of SVs

We applied the program DFE-α v2.15 to estimate the distribution of fitness effects (DFE) and the proportion of adaptive variants (α) for non-synonymous SNPs, DELs, DUPs, INVs, TRAs, and MEIs^89,90^. In these analyses, we used information from synonymous SNPs as the neutral reference, based on the unfolded SFS. First, we fitted a demographic model to the SFS for neutral sites using maximum likelihood (ML). We chose a two-epoch demographic model that allows a single step change in population size from *N*_1_ to *N*_2_*t*_2_ generations in the past^89^. We performed multiple ML searches, each with a different starting point, and treated the parameter values that produced the highest log-likelihood as the ML estimates of the demographic parameters. Next, given the estimated parameters of the demographic model, we inferred the DFE by fitting a γ distribution to the SFS for the selected sites. As above, we carried out multiple searches with different starting values for *β* and *s*, where *β* is the shape parameter of the gamma distribution and *s* is the mean fitness effect of variants. The ML estimates of the DFE parameters and the observed divergence at the selected and neutral sites were then used to estimate the proportion of substitutions (α) that have been fixed by positive selection^90^. We obtained 95% confidence intervals (CIs) for the parameter estimates by analyzing 100 bootstrap replicates of SFS and divergence data sets, which were generated by randomly sampling genes. Following the findings of^91^, we used high-quality data from two North American wild *Vitis* species as outgroup^91^ to infer the ancestral state of variants. We note, however, that the inference of the ancestral state of SVs are likely to be inaccurate, because the genetic divergence between the wild *Vitis* species and Char04 complicated the mapping process. We therefore also used the folded SFS to estimate the DFE and α, using polyDFE v2.0^92^. The results were similar, so we presented the polyDFE results with 95% CIs obtained from the inferred discretized DFEs from 100 bootstrap data sets.

### SVs and sex determination

*F*_ST_ values for both SNPs and SVs exhibited clear outlier peaks in the sex determination region (Figure 3). The SNPs of the sex determination region were phased and imputed based on a genetic map^93^ using Shapeit v2.12^94^, following the study of^15^. To examine relationships among different sex haplotypes, we built Maximum Likelihood (ML) trees from SNPs within the region. ML trees were based on 10,000 bootstrap replicates, as implemented in MEGAX^95^. We built trees for the two regions, corresponding to the peaks of SNP divergence^15^. We reasoned that the true SD region should cluster by gender, which was true for the first peak of the SD region but not the second (Figure S10). We therefore concluded that the first peak, defined as the region between 4.90 Mb and 5.04Mb on chromosome 2 of the PN40024 assembly, represents the SD region. BEAST v1.8.0^96^was applied to calculate genetic divergence, based on a tree with a relaxed molecular clock. After a burn-in of 100,000 steps, data were collected once every 1,000 steps from 10 million MCMC cycles, The divergence time between haplotypes was bases on a genome-wide divergence time of 46.9 million years ago between M. *rotundifolia* and *Vitis* species^97^.

The boundaries of the sex determination region were determined by mapping the coding sequences (CDS) of the chr02:4840000 – 498000 region from PN40024 12X.v2^33^ to the Char04 and Cab08 references. For both Chardonnay and Cabernet Sauvignon haplotypes, gene models were refined by mapping all the CDS identified in the four haplotypes onto Char04 and Cab08 genome assemblies, separately, using GMAP v.2015-11-20 with default parameters^98^.

We analyzed gene expression data from the three grape flower genders. Raw sequencing data were obtained from the Short Read Archive (SRP041212). Reads were first trimmed using Trimmomatic v.0.36^99^ with the options: LEADING:3 TRAILING:3 SLIDINGWINDOW:10:20 MINLEN:20. High-quality reads were mapped onto the primary and haplotig genome assemblies of Char04 and Cab08^17^ separately, using HISAT2 v.2.0.5^100^ with the following options: --end-to-end --sensitive --no-unal. The Bioconductor package GenomicAlignments v.1.12.1^101^ was used to extract counts of uniquely mapped reads (Q > 20). Mapped reads were then normalized by millions of mapped reads per library (RPM). Differential expression analysis across flower genders (i.e. Male vs. Female, Male vs. Hermaphrodite, Female vs. Hermaphrodite) was performed using the Bioconductor package DESeq2 v1.16.1^102^ using samples of the last two flower growth stages as replicates to allow enough statistical power. These same data were analyzed previously using the same methods, based on mapping to the PN40024 reference^15^. The previous work found a tendency toward female biased expression of genes in the sex region. However, in the current analyses the genes that differ in expression in the sex-determination tend to show male-biased expression. The differences between studies reflect mapping biases between the presumed female haplotype in the PN40024^32^ and the H haplotype in the Char04 reference. For these reasons, we consider the gene expression analyses to be a tool to help identify interesting candidate loci, but caution that additional studies of sex biased expression are merited.

### SVs and berry color

We compared genomes of two cultivars with dark blue berries (PN and Cab08) with two cultivars with light green berries (Char04 and Sultanina) using pairwise whole-genome alignments and called SVs using the MUMmer4 pipeline. Dot plots were generated using mumplot from (mumplot -l 100 -c 1000 -d 10 -banded -D 5) for chromosome 2 where the berry color QTL located. For Char04 and Cab08, we verified the SV calls using the Sniffles pipeline^18^ after mapping SMRT reads onto the PN40024 reference genome using both the Minimap2^77^ and NGMLR^18^. We also zoomed in on this region for SV calls for the population samples to investigate the potential association of SVs, gene expression and the berry color in different cultivars.

To identify whether other green berry accessions housed large inversions that include the berry color genes, we determined the orientation of the rearranged chromosome fragments and putative breakpoints from bam files of discordant PE reads and split reads (SP) after mapping short-reads to the PN40024 genome V2.0^33^. Reads were mapped using the BWA-MEM algorithm in bwa-0.7.12^76^. The discordant reads and split reads were extracted using samtools v1.9^55^ and LUMPY v0.2.13^81^. To select breakpoints distinguishing genomes of red- and white-berry cultivars, the discordant, the splitter, and the original bam files were inspected visually using IGV v2.2^103^.

To detect potentially hemizygous regions on chromosome 2, we calculated runs of homozygosity (ROH) for each sample using the software PLINK v1.9^86^ with the following options: --homozyg-window-het 0 --homozyg-snp 41 --homozyg-window-snp 41 --homozyg-window-missing 0 --homozyg-window-threshold 0.05 --homozyg-kb 500 --homozyg-density 5000 --homozyg-gap 1000. CNV analyses were conducted in cnv-seq^104^ using the neighboring grapevines with green and dark blue berry colors with bam file of the former as test while bam file of the later as a reference. The log2 values of the adjusted copy number ratio were plotted in R.

Gene expression analyses of the berry color region utilized the raw RNA-seq data from SRA: SRP049306-SRP049307^43^. The data were generated from berries sampled during berry development at four stages, including two before and two after veraison, from 10 Italian varieties (5 red and 5 white). RNA-seq data were mapped onto the Char04 reference and analyzed as described in the previous section. Differential gene expression analysis was performed for each berry growth stage, separately, by comparing samples from red cultivars with berries from with varieties. Genes presented an adjusted *P*-value ≤ 0.05 between red and white cultivars were considered as significantly expressed. Gene expression analyses focused on the 173 genes in the Char04 chromosome 2 inversion.

## END NOTES

### Acknowledgements

We are grateful for the technical assistance of Rosa Figueroa-Balderas, the services of the Genomics High Throughput Facility at UC Irvine, and the comments of A. Muyle, D. Seymour, D. Koenig, T. Batarseh, G. Martin and J. Ross-Ibarra. This work was supported by seed funding from UC Irvine, NSF grants 1542703 to BSG, NSF grant #1741627 to BSG and DC, and support to DC by J. Lohr Vineyards and Wines, E. & J. Gallo Winery, and the Louis P. Martini Endowment in Viticulture.

### Author Contributions

YZ, AM, ES, MM, DC and BG wrote the manuscript. YZ, AM, MM and YL performed analyses. TB provided data. DC and BG supervised and guided the research.

