## Supplementary file for "Structural variants, hemizygosity and clonal propagation in grapevines"

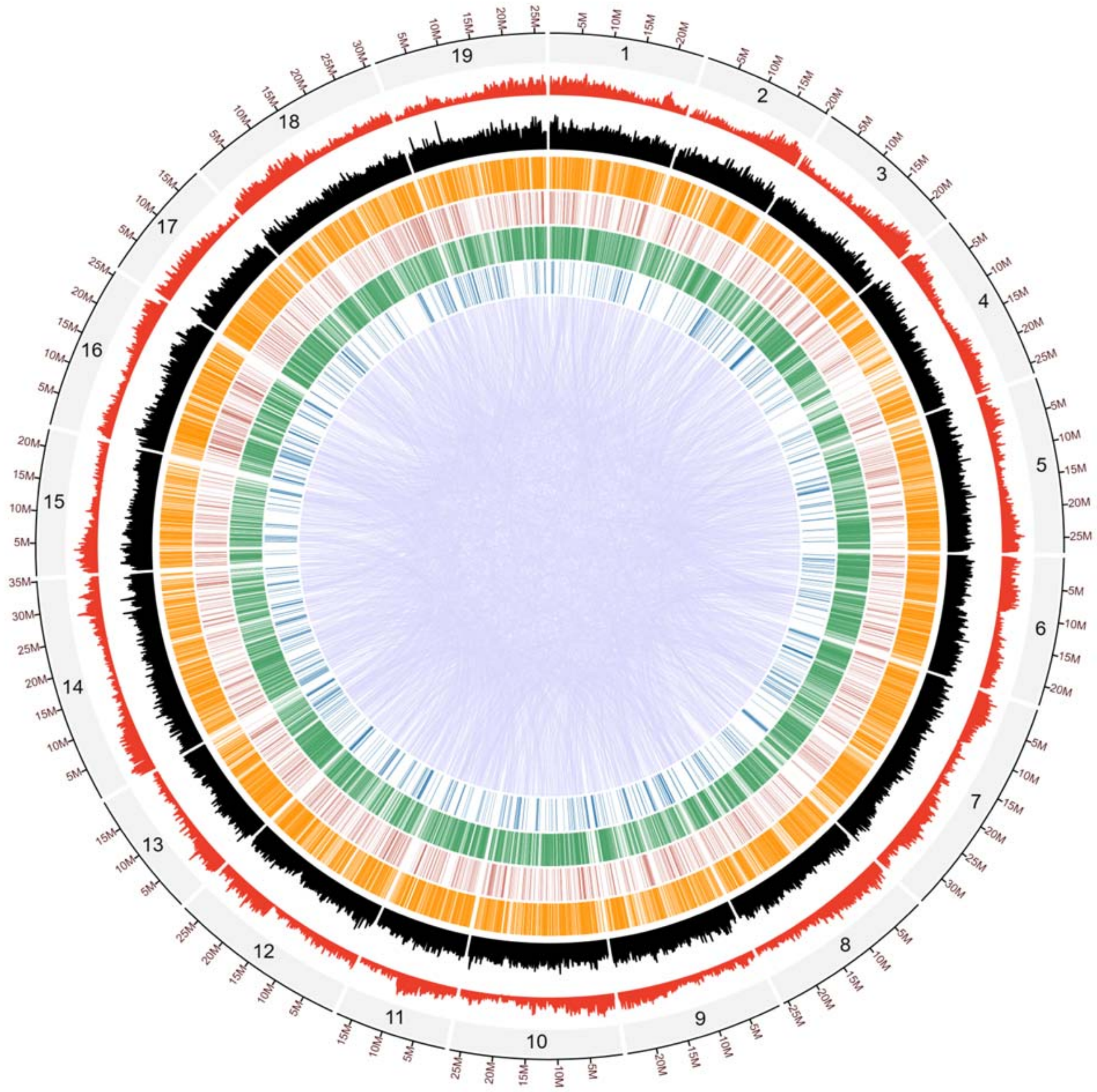


**Fig. S1**

Structural heterozygosity between Char04 and Cab08. The circle plot reports SVs detected between Char04 and Cab08 from mapping Cab08 SMRT reads to the Char04 reference. The outermost circle denotes the number and size of chromosomes (gray), followed by gene density (red), TE density (black), deletion SVs (orange), duplications (dark red), small insertions (green), inversions (blue) and with translocations represented in the middle of the circle in purple.

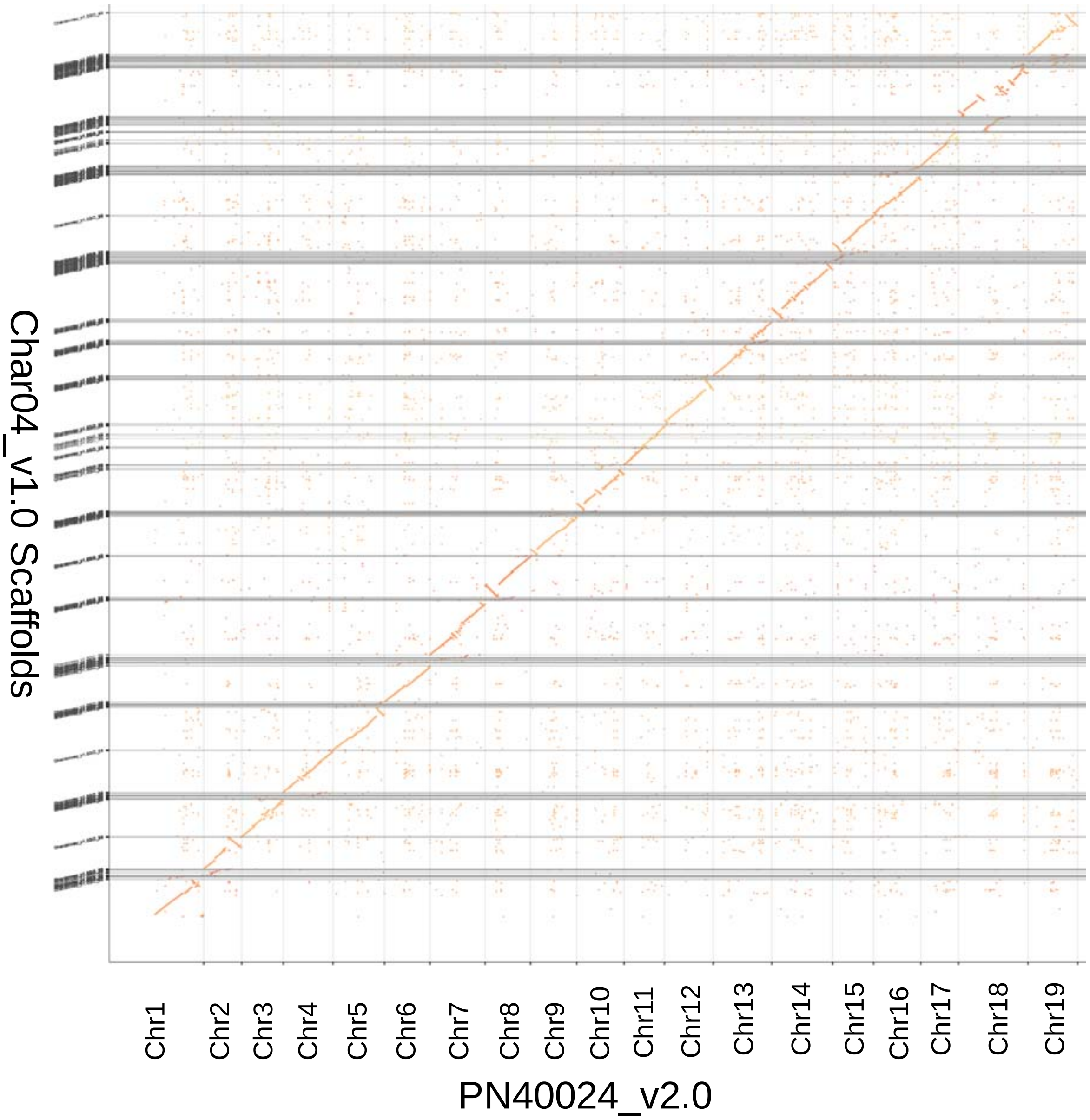


**Fig. S2.**

A dot plot demonstrates high colinearity between the PN40024 assembly and the Char04 assembly, but also with a number of discrete inversions, such as the one encompassing the berry color region on chromosome 2.

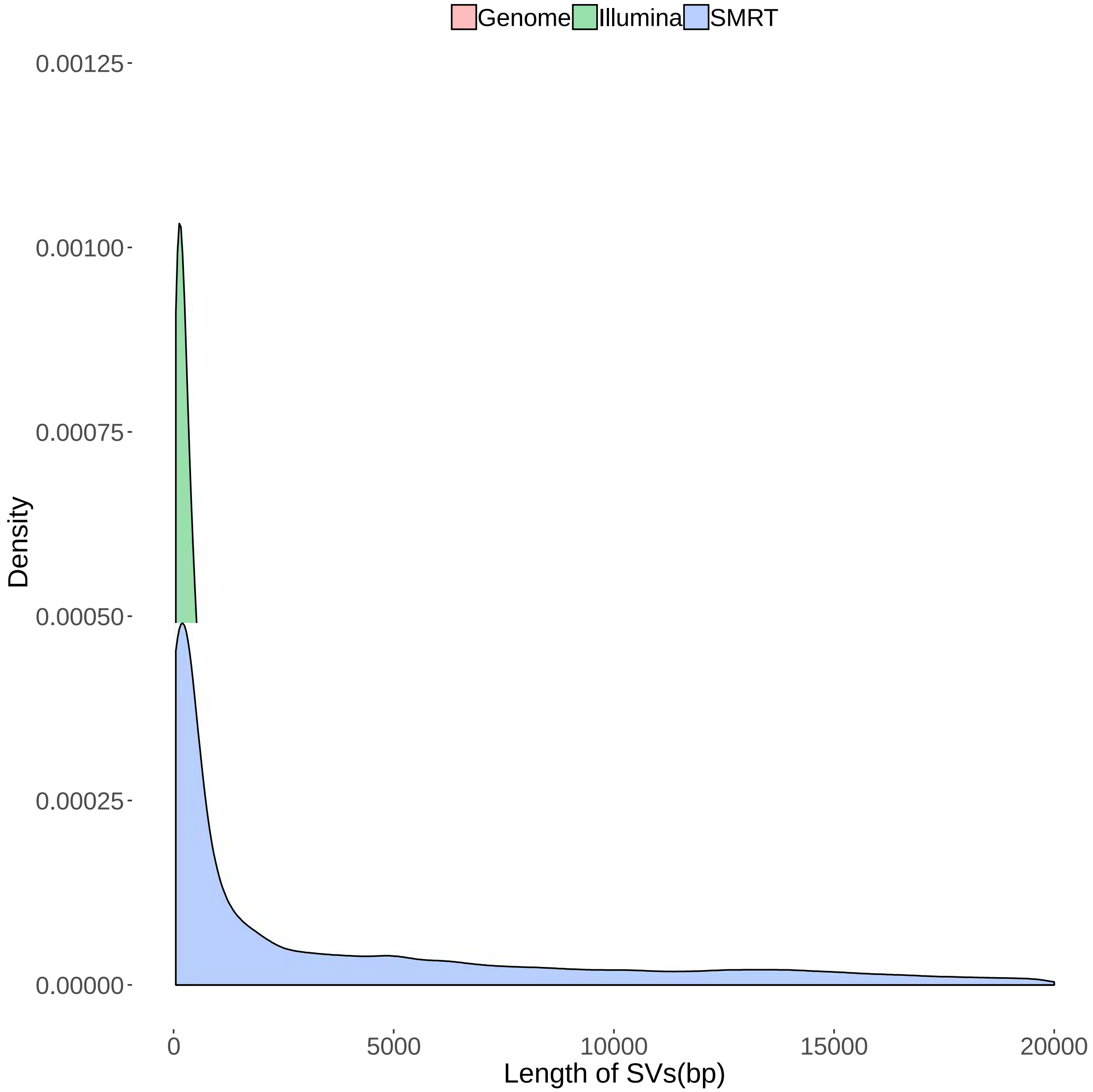


**Fig. S3**

The length distributions of SVs between Cab08 and Char04 detected by genome alignment, long reads (PacBio), and Illumina paired short reads.

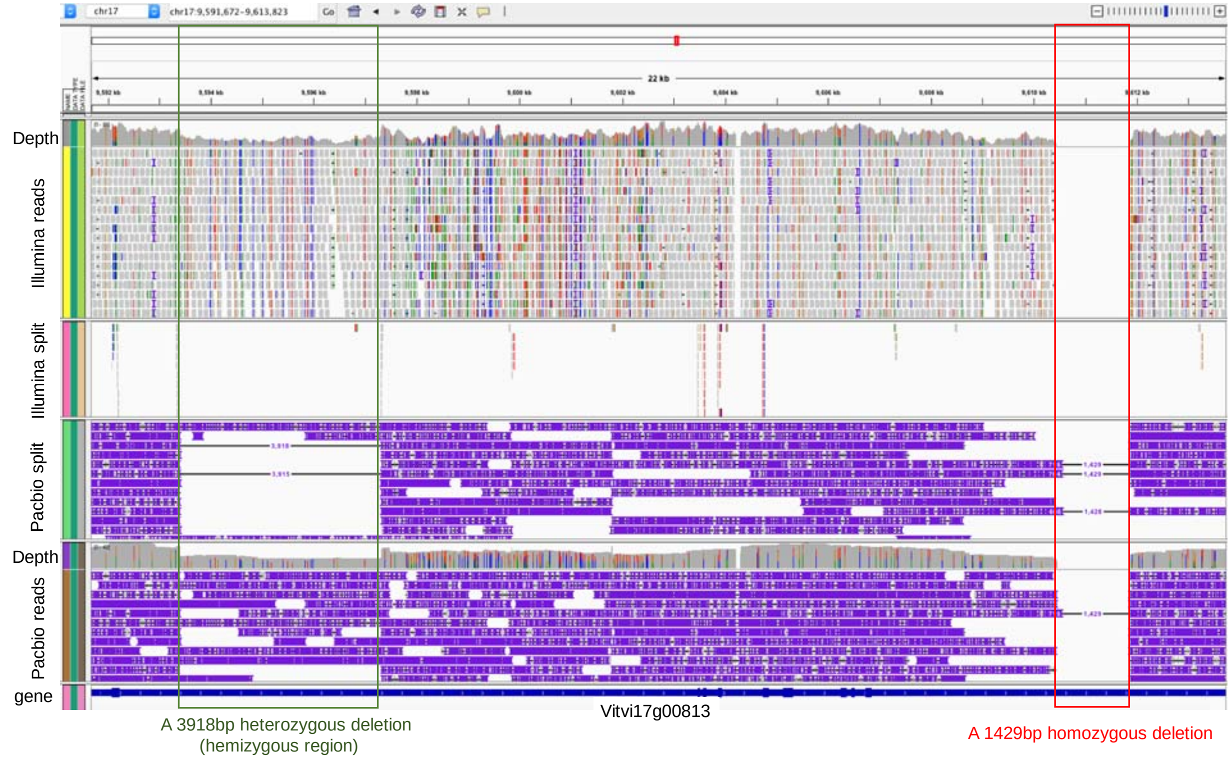


**Figure S4:** A demonstration of SV inferences, based on mapping SMRT long reads and short-reads from Cab08 to Char04. This region houses two unlinked SVs, a 3,918 bp heterozygous deletion and a 1,429 bp homozygous deletion within a single gene.

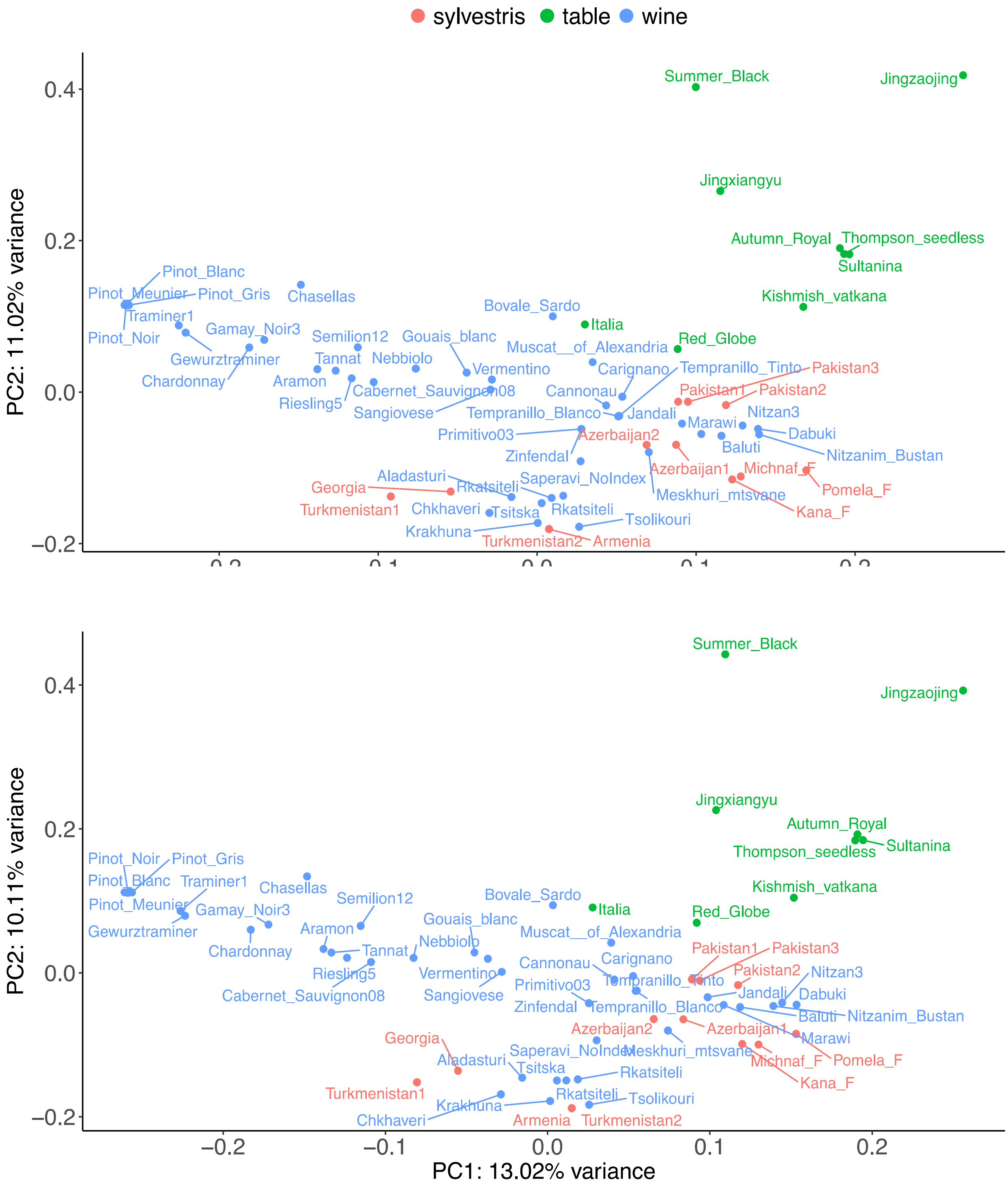


**Fig. S5.**

Principal component analyses (PCA) of the entire sample of wild and cultivated *V. vinifera* grapevines for SNPs (top) and SVs (bottom). For both graphs, sites with minor allele frequencies ≤ 0.05 and ≥ 20% missing data were removed.

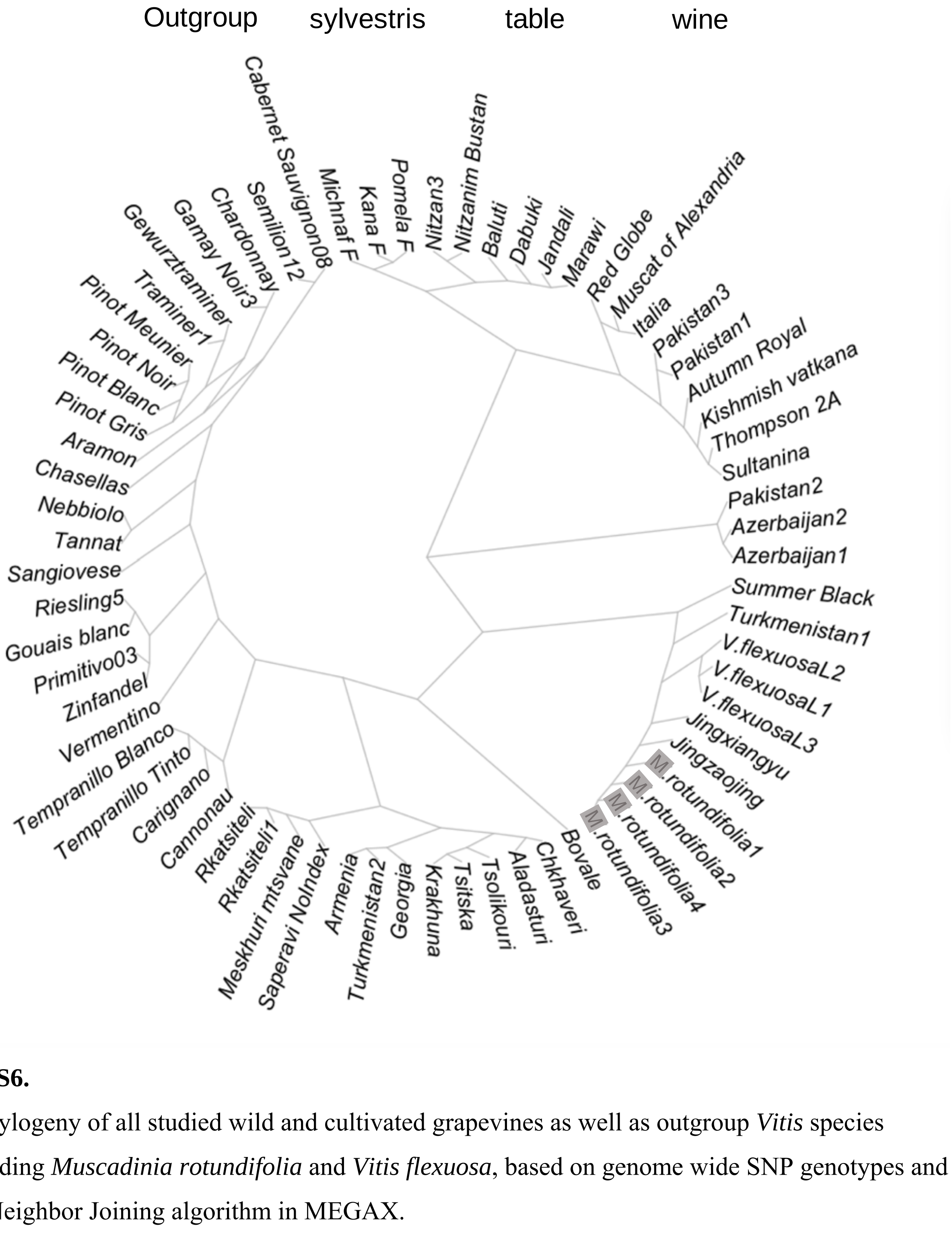


**Fig. S6.**

A phylogeny of all studied wild and cultivated grapevines as well as outgroup *Vitis* species including *Muscadinia rotundifolia* and *Vitis flexuosa*, based on genome wide SNP genotypes and the Neighbor Joining algorithm in MEGAX.

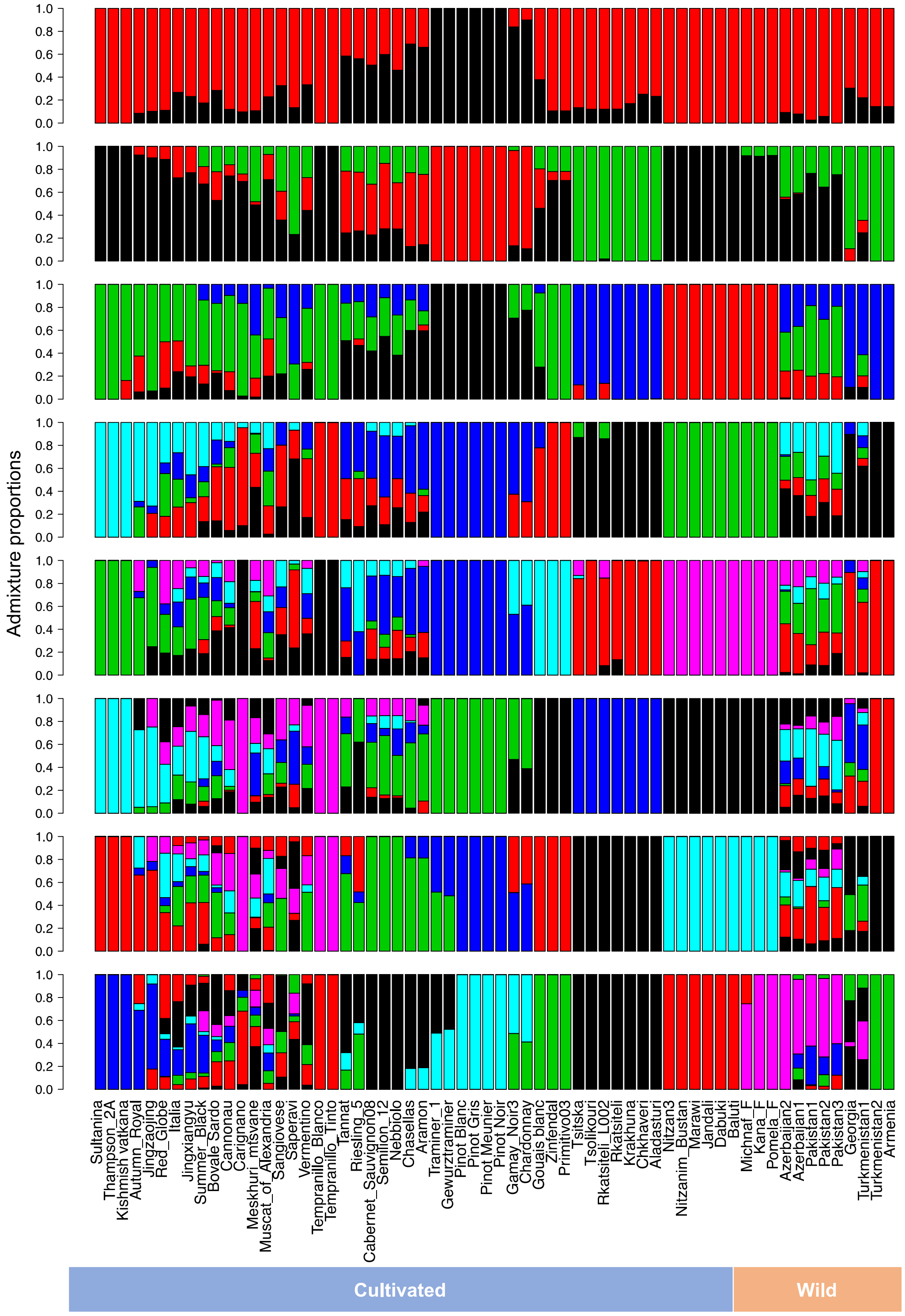


**Fig. S7.**

Population structure analyses of wild and cultivated grapevines. Structure plots are based on $K$=2 (top) sequentially to $K$=9 (bottom). $K$ = 8 is the most likely number of clusters to explain the data. The Structure analysis was used to help guide down-sampling to $n$=12 for Figure 2.

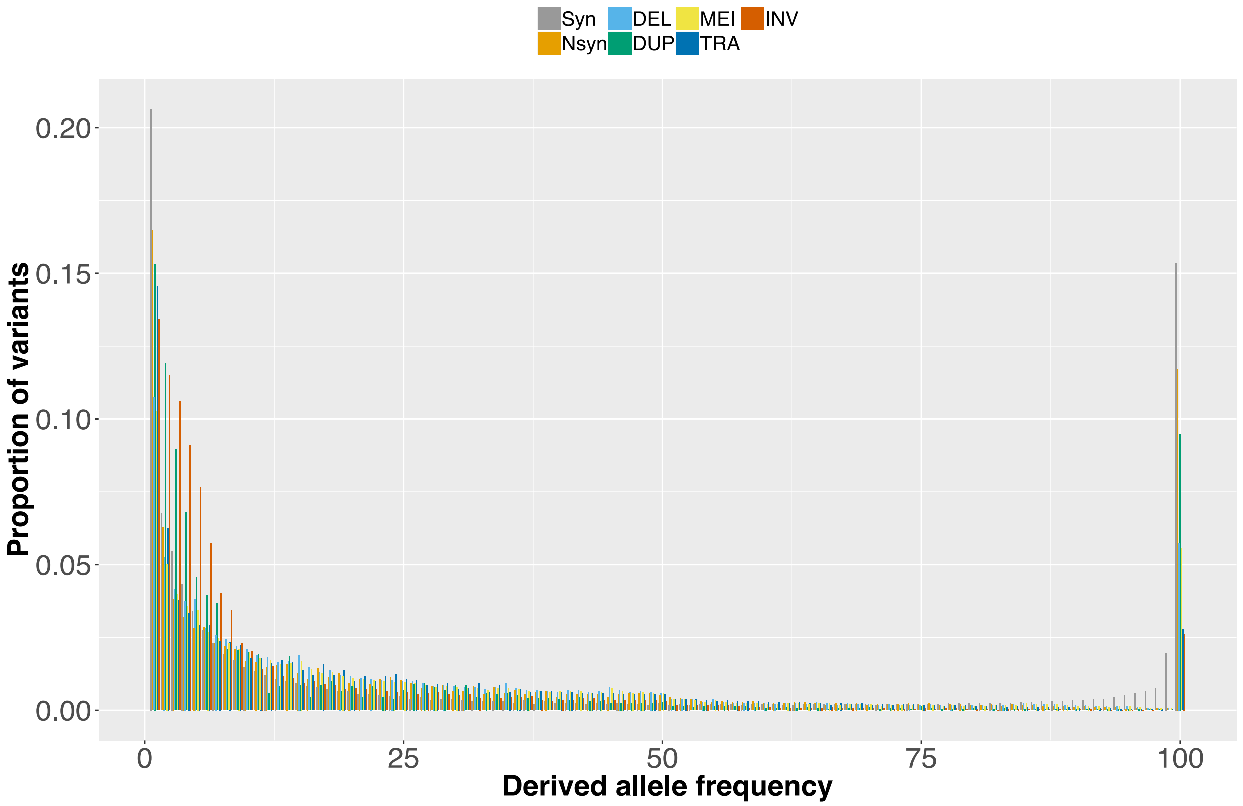


**Fig. S8.**

The unfolded SFS for all 50 cultivars in this study. The discrete drop in frequencies at 50% allele frequency is evident in this sample, as it was in the cultivated sample in Figure 2. The SV categories include synonymous SNPs (Syn), nonsynonymous SNPs (Nsyn), deletions (DEL), duplications (DUP), TE insertions (MEI), translocations (TRA) and inversions (INV).

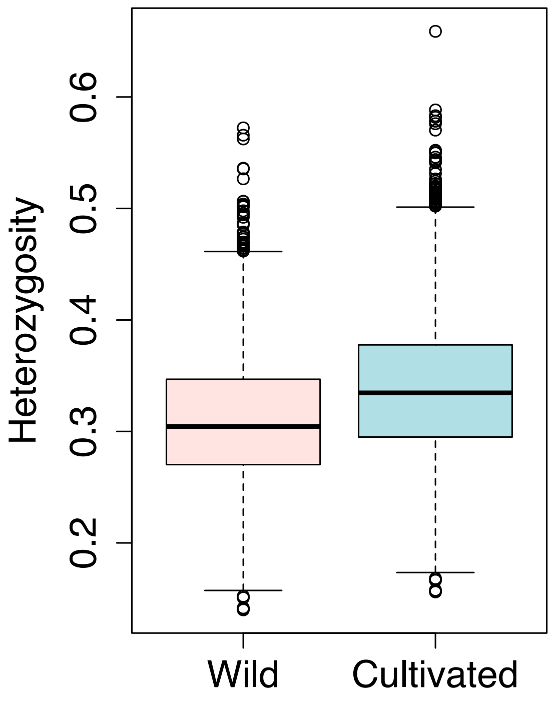


**Fig. S9.**

The observed heterozygosity ($H_o$) within wild ($H_{o\_wild} = 0.304 \pm 0.059$) and cultivated ($H_{o\_cult} = 0.339 \pm 0.0625$) grapevines, based on 10,972,223 SNPs and 20 Kb, non-overlapping windows. Cultivated *sativa* is 11.31% more heterozygous, on average, than the individuals in our wild sample of *sylvestris* accessions.

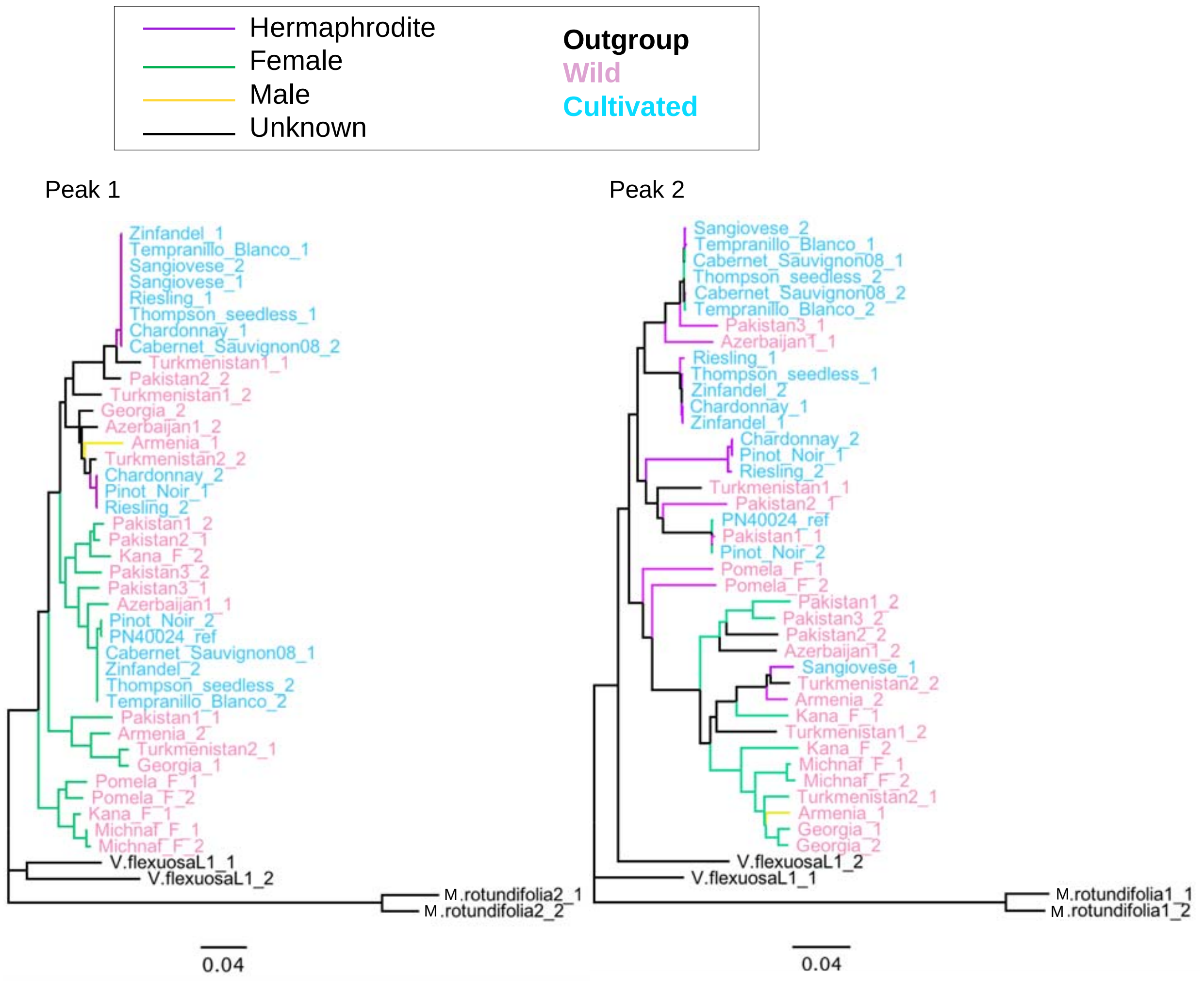


**Fig. S10.**

Phylogeny of haplotypes based on two peaks of genomic differentiation underlying sex determination detected by (*2*). The first peak (left) contains the sex region, and the sequences separate according to sex haplotype. The second peak (right) demonstrates a different history that is not (apparently) correlated with sex. These two phylogenies imply that the second peak does not correspond to sex determination and that there is recombination between the regions defined by the two peaks. The function of the second peak, and the reason for large divergence between wild and cultivated (Figure 3), remains unclear.

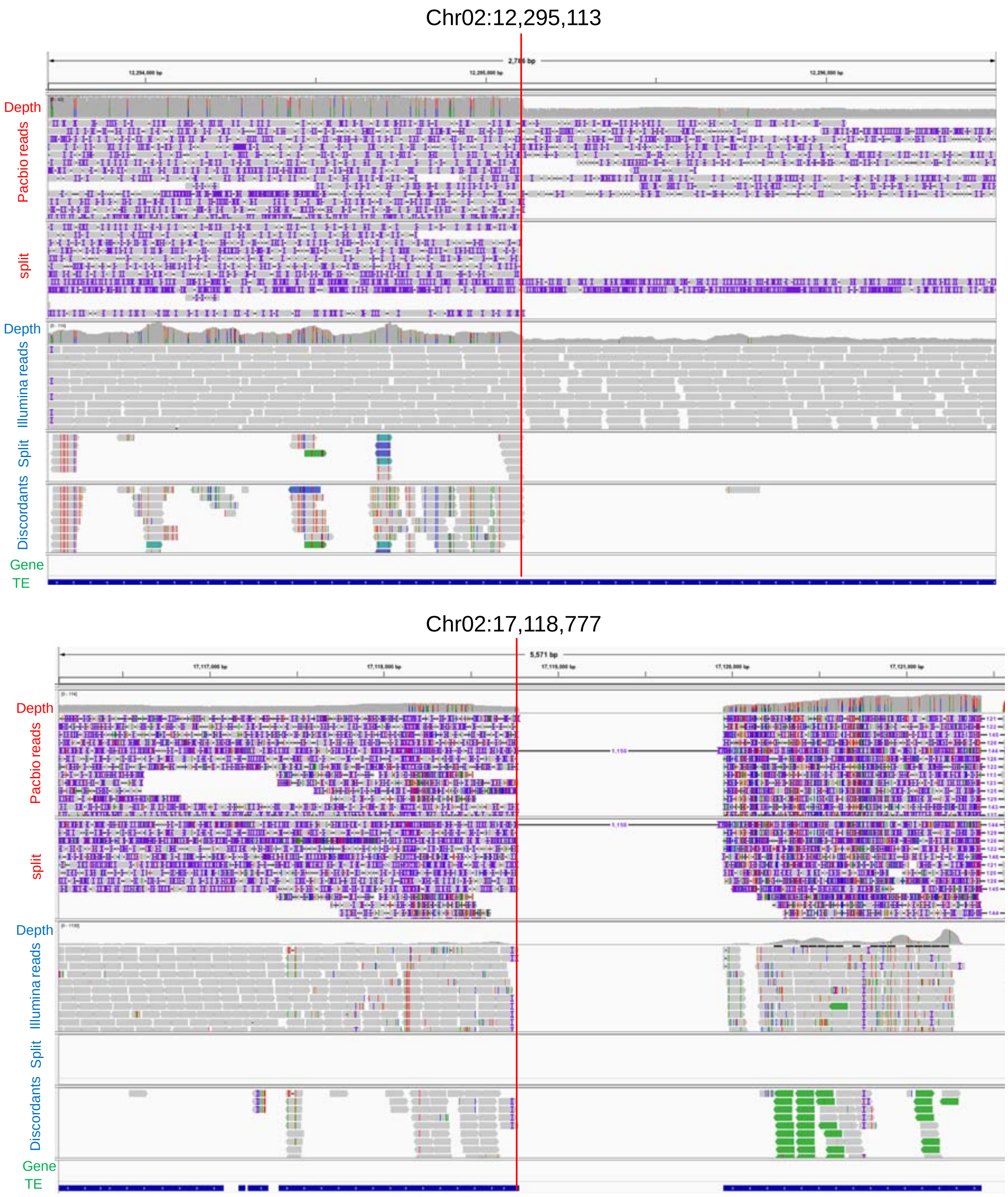


**Fig. S11**. The Integrated Genomve Viewer view of a 4.82Mb inversion (chr02: 12,295,113bp-17,118,777bp) in Char04 mapped to PN40024 reference based on Pacbio long reads and Illumina short-reads. Both the read depth and heterozygosity (vertical colored lines in the grey coverage plots) support that the inverted region is in a hemizygous state. Breakpoints on both sides locate in TEs.

**Table S1.**

Comparison of TE components of Char04, Cab08, and PN40024 genome

| | | PN40024 V3 | | | Cabernet Sauvignon | | | Chardonnay | | |
|---|---|---|---|---|---|---|---|---|---|---|
| | | *Chromosomes (with ChrUN)* | | | *Primary* | | | *Primary* | | |
| Class | | Count | Cumulative length (bp) | Assembly % | Count | Cumulative length (bp) | Assembly % | Count | Cumulative length (bp) | Assembly % |
| *LINEs* | *L1* | 3,850 | 1,324,786 | 0.3% | 16,656 | 23,822,207 | 4.0% | 25,246 | 25,084,055 | 4.2% |
| | *L2* | 40 | 24,557 | 0.0% | 6 | 14,601 | 0.0% | 34 | 23,455 | 0.0% |
| | *Penelope* | 201 | 30,534 | 0.0% | 93 | 37,981 | 0.0% | 234 | 29,909 | 0.0% |
| | *R2* | 14 | 5,270 | 0.0% | 2 | 1,402 | 0.0% | 22 | 8,142 | 0.0% |
| | *RTE-BovB* | 87 | 14,060 | 0.0% | 70 | 13,492 | 0.0% | 86 | 14,290 | 0.0% |
| *SINEs* | | 12 | 5,025 | 0.0% | 52 | 15,979 | 0.0% | 22 | 6,408 | 0.0% |
| *LTRs* | *Caulimovirus* | 4,027 | 2,441,413 | 0.5% | 2,202 | 6,243,275 | 1.1% | 5,384 | 3,578,875 | 0.6% |
| | *Copia* | 42,521 | 18,533,372 | 3.9% | 38,091 | 64,564,145 | 10.9% | 86,345 | 66,745,378 | 11.2% |
| | *ERV1* | 94 | 38,460 | 0.0% | 67 | 35,843 | 0.0% | 127 | 47,460 | 0.0% |
| | *Gypsy* | 85,844 | 57,680,169 | 12.3% | 50,360 | 136,034,576 | 23.0% | 130,121 | 100,285,972 | 16.9% |
| | *Ngaro* | 34 | 15,812 | 0.0% | 40 | 15,955 | 0.0% | 50 | 24,014 | 0.0% |
| | *Pao* | 44 | 35,282 | 0.0% | 58 | 50,084 | 0.0% | 43 | 34,194 | 0.0% |
| | *RC* | - | - | 0.0% | 162 | 136,667 | 0.0% | -- | -- | |
| | *Helitron* | 240 | 81,361 | 0.0% | 759 | 564,628 | 0.1% | 1,103 | 597,973 | 0.1% |
| | *Other LTR* | 20,418 | 10,227,279 | 2.2% | 9,469 | 11,758,127 | 2.0% | 25,282 | 12,373,164 | 2.1% |
| *TRIMs* | | | | 0.0% | 5,276 | 3,118,302 | 0.5% | 17,265 | 4,860,759 | 0.8% |
| *DNA Transposons* | *CMC-Chapaev* | 223 | 64,038 | 0.0% | 45 | 124,641 | 0.0% | 278 | 67,224 | 0.0% |
| | *CMC-EnSpm* | 13,965 | 4,108,835 | 0.9% | 8,746 | 9,266,212 | 1.6% | 18,785 | 5,758,505 | 1.0% |
| | *CMC-Transib* | 299 | 50,228 | 0.0% | 96 | 37,953 | 0.0% | 386 | 59,857 | 0.0% |
| | *En-Spm* | 337 | 207,713 | 0.0% | 190 | 232,586 | 0.0% | 456 | 274,196 | 0.0% |
| | *Harbinger* | 11 | 6,014 | 0.0% | 1,811 | 1,421,983 | 0.2% | 19 | 6,409 | 0.0% |
| | *hAT* | 8,731 | 2,536,441 | 0.5% | 7,192 | 8,378,344 | 1.4% | 15,728 | 8,513,667 | 1.4% |
| | *Tc1* | - | - | 0.0% | 35 | 24,041 | 0.0% | | | |
| | *ULE-MuDR, Mul* | 23,452 | 9,057,320 | 1.9% | 16,070 | 19,511,700 | 3.3% | 37,382 | 20,633,755 | 3.5% |
| | *PIF-Harbinger* | 29,392 | 6,162,660 | 1.3% | 17,100 | 7,061,177 | 1.2% | 41,266 | 9,732,500 | 1.6% |
| | *TcMar-Fot1* | 42 | 10,011 | 0.0% | 14 | 6,701 | 0.0% | 52 | 10,549 | 0.0% |
| | *TcMar-Pogo* | 175 | 37,769 | 0.0% | 137 | 45,073 | 0.0% | 238 | 48,802 | 0.0% |
| | *Other DNA* | 1,995 | 653,842 | 0.1% | 2,202 | 1,153,276 | 0.2% | 2,715 | 913,405 | 0.2% |
| *Others* | | 107,789 | 107,675,226 | 22.9% | 15,484 | 8,720,028 | 1.5% | 33,405 | 21,761,307 | 3.7% |
| ***Total interspersed Repeats*** | | ***343,837*** | ***221,027,477*** | ***47.0%*** | ***192,485*** | ***302,410,979*** | ***51.1%*** | ***442,074*** | ***281,494,224*** | ***47.3%*** |

**Table S2.**

The number of variants detected by genome alignment, Pacbio long reads, and Illumina short reads.

| Variant type | Illumina short reads | | Pacbio long reads | Genome alignment |
| --- | --- | --- | --- | --- |
| | Heterozygous | Homozygous | | |
| Char04 to Char04 | | | | |
| SNPs & indels | 1,179,253 | 2,557 | 992,226 | |
| DEL | 9,440 | 82 | 8,302 | |
| DUP | 722 | 10 | 891 | |
| INV | 140 | 5 | 395 | |
| TRA | 672 | 12 | 1,355 | |
| MEI | 4,348 | 31 | 7,772 | |
| CNV | | | 273 | |
| Cab08 to Char04 | | | | |
| SNPs & indels | 1,166,351 | 349,440 | 998,147 | |
| DEL | 19,065 | 3,817 | 24,138 | 28,089 |
| DUP | 2,698 | 633 | 3,544 | 9,267 |
| INV | 627 | 44 | 1,601 | 1,513 |
| TRA | 2,211 | 318 | 5,744 | 3,786 |
| MEI | 12,705 | 2,253 | 21,722 | 10,043 |
| CNV | | | 3,164 | 1,254 |

**Table S3.**

Population samples of wild and cultivated grapevines used in this study

| Accessions | Clone | Country | Seed | Berry color | Group | SRA | Reference[4] |
|---|---|---|---|---|---|---|---|
| Aladasturi | | Georgia | 1 | db | w | | 1 |
| Aramon | | France | 1 | db | w | SRR5627800 | 2 |
| Armenia | | | 1 | | wild | SRR5627784 | 2 |
| Autumn Royal | | CA | 0 | db | t | SRR354199 | 3 |
| Azerbaijan1 | | | 1 | | wild | SRR5627787 | 2 |
| Azerbaijan2 | | | 1 | | wild | SRR5627790 | 2 |
| Cabernet_Sauvignon08 | 8 | CA | 1 | db | w | SRR5627795 | 2 |
| Chardonnay | 4 | CA | 1 | g | w | SRR5627799 | 2 |
| Chasellas | | INRA | 1 | g | w | | 1 |
| Gewurztraminer | | Italy | 1 | pink-red | w | ERR514999 | 4 |
| Gamay_Noir | 3 | | 1 | db | w | SRR5627798 | 2 |
| Georgia | | | 1 | | wild | SRR5627789 | 2 |
| Gouais blanc | | INRA | 1 | g | w | | 1 |
| Italia | | | 1 | g | t | SRR354198 | 3 |
| Krakhuna | | Georgia | 1 | g | w | | 1 |
| Muscat of Alexandria | | North Africa | 1 | g | w | SRR5627781 | 2 |
| Chkhaveri | N3 | Georgia | 1 | db | w | | 5 |
| Meskhuri mtsvane | N4 | Georgia | 1 | g | w | | 5 |
| Pakistan1 | | | 1 | | wild | SRR5627785 | 2 |
| Pakistan2 | | | 1 | | wild | SRR5627792 | 2 |
| Pakistan3 | | | 1 | | wild | SRR5627791 | 2 |
| Primitivo03 | | Italy | 1 | db | w | SRR5627796 | 2 |
| Red_Globe | | | 1 | pink-red | t | SRR354201 | 3 |

| | | | | | | | |
|---|---|---|---|---|---|---|---|
| Riesling | | INRA | 1 | g | w | SRR5627794 | 2 |
| Rkatsiteli | | Georgia | 1 | g | w | | 5 |
| Rkatsiteli_1 | | Georgia | 1 | g | w | | 1 |
| Saperavi | | Georgia | 1 | db | w | | 5 |
| Semilion | 12 | France | 1 | g | w | SRR5627793 | 2 |
| Nitzanim_Bustan | | Israel | 1 | g | w | SRR3496931 | 1 |
| Pomela_wild_female | | Israel | 1 | db | wild | SRR3497168 | 6 |
| Nitzan3 | | Israel | 1 | g | w | SRR3508973 | 6 |
| Kana_wild_female | | Israel | 1 | db | wild | SRR3509067 | 6 |
| Pinot Gris | | France | 1 | pink | w | SRR3509358 | 7 |
| Pinot Blanc | | France | 1 | g | w | SRR3509362 | 7 |
| Pinot Meunier | | France | 1 | db | w | SRR3509365 | 7 |
| Michnaf_wild_female | | Israel | 1 | | wild | SRR3509717 | 6 |
| Baluti | | Israel | 1 | g | w | SRR3509718 | 6 |
| Marawi | | Israel | 1 | g | w | SRR3509720 | 6 |
| Dabuki | | Israel | 1 | g | w | SRR3528127 | 6 |
| Jandali | | Israel | 1 | g | w | SRR3528198 | 6 |
| Pinot Noir | | | 1 | db | w | SRR3990782 | 2 |
| Nebbiolo_CVT423 | Picoutenter | Italy | 1 | db | w | SRR5626750 | 8 |
| Kishmish vatkana | | Uzbekistan | 0 | db | t | SRR5712111 | 9 |
| Cannonau | | Sardinia | 1 | db | w | SRR5803836 | 10 |
| Bovale Sardo | | Sardinia | 1 | db | w | SRR5803837 | 10 |
| Vermentino | | Sardinia | 1 | g | w | SRR5803838 | 10 |
| Carignano | | Sardinia | 1 | db | w | SRR5803839 | 10 |
| Muscares_Carlos_B0017EY | | France:Colmar | 1 | | outgroup | SRR6729327 | 11 |
| Muscares_Regale_B00174T | | France:Colmar | 1 | | outgroup | SRR6729330 | 11 |
| Muscares_Regale_B00FFX1 | | France:Colmar | 1 | | outgroup | SRR6729331 | 11 |
| Jingxiangyu | | China | 0 | g | t | SRR769818 | 12 |
| Jingzaojing | | China | 0 | g | t | SRR769819 | 12 |

| | | | | | | | |
|---|---|---|---|---|---|---|---|
| Sangiovese | | Italy | 1 | db | w | SRR5506711 | 13 |
| Tannat | UY11 | Uruguay | | | w | SRR863618 | 14 |
| sultanina | | Chile | 0 | g | t | SRR931841-46 | 15 |
| Summer_Black | | China | 0 | b | t | SRR2967092 | 3 |
| Tempranillo Blanco | | Spain | 1 | g | w | SRR2895165 | 16 |
| Tempranillo Tinto | | Spain | 1 | db | w | SRR2895164 | 16 |
| Thompson Seedless | 2A | Davis | 0 | g | t | SRR5627782 | 2 |
| Traminer | 1 | Italy | 1 | pink-red | w | SRR5627802 | 2 |
| Tsitska | | Georgia | 1 | g | w | | 1 |
| Tsolikouri | | Georgia | 1 | g | w | | 1 |
| Turkmenistan1 | | | 1 | | wild | SRR5627783 | 2 |
| Turkmenistan2 | | | 1 | | wild | SRR5627786 | 2 |
| *Muscadinia_rotundifolia1* | | | 1 | | outgroup | SRR5627788 | 2 |
| Zinfandel_03 | | CA | 1 | db | w | SRR5627801 | 2 |
| *V.flexuosa*_L-10 | | | 1 | | Vitis | SRR6877426 | 17 |
| *V.flexuosa*_L-200 | | | 1 | | Vitis | SRR6877427 | 17 |
| *V.flexuosa*_L-2 | | | 1 | | Vitis | SRR6877428 | 17 |

[1] Seed (1); seedless (0);
[2] Berry color: green (g); dark blue (db);
[3] Group: wine (w); table (t)
[4] References to Table S3:

1. Tabidze, Vazha, Grigol Baramidze, Ia Pipia, Mari Gogniashvili, Levan Ujmajuridze, Tengiz Beridze, Alvaro G. Hernandez, and Barbara Schaal. 2014. “The Complete Chloroplast DNA Sequence of Eleven Grape Cultivars. Simultaneous Resequencing Methodology.” J Int Sci Vigne Vin 48:99–109.
2. Zhou, Yongfeng, Mélanie Massonnet, Jaleal S. Sanjak, Dario Cantu, and Brandon S. Gaut. 2017. “Evolutionary Genomics of Grape (Vitis Vinifera Ssp. Vinifera) Domestication.” Proceedings of the National Academy of Sciences of the United States of America 114 (44): 11715–20.
3. Cardone, Maria Francesca, Pietro D’Addabbo, Can Alkan, Carlo Bergamini, Claudia Rita Catacchio, Fabio Anaclerio, Giorgia Chiatante, et al. 2016. “Inter-Varietal Structural Variation in Grapevine Genomes.” The Plant Journal: For Cell and Molecular

Biology 88 (4): 648–61.
4. Leonardelli, Lorena, Alessandro Cestaro, Carmen Maria Livi, Patrice This, and Claudio Moser. 2013. “VerySNP: VCF Features to Train SVM in Crop SNP Detection.” Conference: 21th Annual Internation Conference on Intelligence Systems for Molecular Biology, 12th European Conference on Computational BiologyAt: Berlin
5. Tabidze V, Pipia I, Gogniashvili M, Kunelauri N, Ujmajuridze L, Pirtskhalava M, Vishnepolsky B, Hernandez AG, Fields CJ, Beridze T (2017) Whole genome comparative analysis of four Georgian grape Cultivars. Mol Genet Genomics, ;292(6):1377-1389
6. Drori, Elyashiv, Oshrit Rahimi, Annarita Marrano, Yakov Henig, Hodaya Brauner, Mali Salmon-Divon, Yishay Netzer, et al. 2017. “Collection and Characterization of Grapevine Genetic Resources (Vitis Vinifera) in the Holy Land, towards the Renewal of Ancient Winemaking Practices.” Scientific Reports 7 (March): 44463.
7. Marroni, Fabio, Davide Scaglione, Sara Pinosio, Alberto Policriti, Mara Miculan, Gabriele Di Gaspero, and Michele Morgante. 2017. “Reduction of Heterozygosity (ROH) as a Method to Detect Mosaic Structural Variation.” Plant Biotechnology Journal 15 (7): 791–93.
8. Gambino, Giorgio, Alessandra Dal Molin, Paolo Boccacci, Andrea Minio, Walter Chitarra, Carla Giuseppina Avanzato, Paola Tononi, et al. 2017. “Whole-Genome Sequencing and SNV Genotyping of ‘Nebbiolo’ (Vitis Vinifera L.) Clones.” Scientific Reports 7 (1): 17294.
9. Canaguier, A., J. Grimplet, G. Di Gaspero, S. Scalabrin, E. Duchêne, N. Choisne, N. Mohellibi, et al. 2017. “A New Version of the Grapevine Reference Genome Assembly (12X.v2) and of Its Annotation (VCost.v3).” Genomics Data 14 (December): 56–62.
10. Mercenaro, Luca, Giovanni Nieddu, Andrea Porceddu, Mario Pezzotti, and Salvatore Camiolo. 2017. “Sequence Polymorphisms and Structural Variations among Four Grapevine (Vitis Vinifera L.) Cultivars Representing Sardinian Agriculture.” Frontiers in Plant Science 8 (July): 1279.
11. PRJNA397021: https://www.ncbi.nlm.nih.gov/bioproject/PRJNA397021
12. PRJNA192798: https://www.ncbi.nlm.nih.gov/bioproject/PRJNA192798
13. Dal Santo, Silvia, Sara Zenoni, Marco Sandri, Gabriella De Lorenzis, Gabriele Magris, Emanuele De Paoli, Gabriele Di Gaspero, et al. 2018. “Grapevine Field Experiments Reveal the Contribution of Genotype, the Influence of Environment and the Effect of Their Interaction (G×E) on the Berry Transcriptome.” The Plant Journal: For Cell and Molecular Biology 93 (6): 1143–59.
14. Da Silva, Cecilia, Gianpiero Zamperin, Alberto Ferrarini, Andrea Minio, Alessandra Dal Molin, Luca Venturini, Genny Buson, et al. 2013. “The High Polyphenol Content of Grapevine Cultivar Tannat Berries Is Conferred Primarily by Genes That Are Not Shared with the Reference Genome.” The Plant Cell 25 (12): 4777–88.
15. Di Genova, Alex, Andrea Miyasaka Almeida, Claudia Muñoz-Espinoza, Paula Vizoso, Dante Travisany, Carol Moraga, Manuel Pinto, Patricio Hinrichsen, Ariel Orellana, and Alejandro Maass. 2014. “Whole Genome Comparison between Table and Wine Grapes Reveals a Comprehensive Catalog of Structural Variants.” BMC Plant Biology 14 (January): 7.
16. Carbonell-Bejerano, Pablo, Carolina Royo, Rafael Torres-Pérez, Jérôme Grimplet, Lucie Fernandez, José Manuel Franco-Zorrilla, Diego Lijavetzky, et al. 2017a. “Catastrophic Unbalanced Genome Rearrangements Cause Somatic Loss of Berry Color in Grapevine.” Plant Physiology 175 (2): 786–801.
17. PRJNA445217: https://www.ncbi.nlm.nih.gov/bioproject/PRJNA445217

**Table S4.**

The number of SNPs (1bp), indels (<50bp) and SVs (≥50bp) used in population analyses. SVs were filtered based on Pacbio SVs calls in Cab08 mapped to Char04.

| Variants | Number of variants |
|---|---|
| SNPs | 15,222,164 |
| Indels | 2,178,370 |
| DEL (deletion) | 238,492 |
| DUP (duplication) | 40,803 |
| MEI (mobile elements insertions) | 181,313 |
| INV (Inversion) | 5,085 |
| TRA (Translocation) | 1,5403 |

**Table S5.**

Fixed SVs between wild and cultivated grapevines and their annotations.

| Type | Chr. | Start | End | geneV3 | Annotation |
|---|---|---|---|---|---|
| DEL | chr02 | 4858421 | 4859731 | | |
| DEL | chr02 | 4909443 | 4910104 | | |
| INS | chr03 | 1590160 | 1590161 | Vitvi03g00126 | NUCLEAR FUSION DEFECTIVE 4-like(NFD4) |
| DEL | chr03 | 10424797 | 10425344 | | |
| DEL | chr03 | 10760472 | 10760533 | | |
| DEL | chr03 | 15495907 | 15495946 | Vitvi03g00132 | pentatricopeptide repeat-containing protein At1g11290 (DOT4) |
| DEL | chr03 | 15542166 | 15542239 | | |
| DEL | chr04 | 15361204 | 15404162 | | |
| DEL | chr04 | 15371981 | 15374475 | Vitvi04g01045 | Vitis vinifera uncharacterized LOC100262060 |
| DEL | chr04 | 15379905 | 15380060 | Vitvi04g01046 | unknown |
| DEL | chr05 | 8742057 | 8742133 | | |
| DUP | chr05 | 8981426 | 9016413 | | |
| DUP | chr05 | 8983782 | 9002151 | Vitvi05g00805 | E3 ubiquitin-protein ligase (UPL1) |
| DEL | chr05 | 9011226 | 9016343 | | |
| DUP | chr05 | 9018177 | 9023809 | | |
| DUP | chr05 | 9023621 | 9023825 | | |
| DEL | chr05 | 9023792 | 9024270 | | |
| DEL | chr05 | 9024335 | 9024923 | | |
| DEL | chr05 | 11257504 | 11257554 | | |
| DEL | chr05 | 22360296 | 22360314 | Vitvi05g00805 | isoleucine--tRNA ligase (OVA2) |
| DEL | chr07 | 12077243 | 12078985 | | |
| INS | chr07 | 19267932 | 19267933 | Vitvi07g01374 | calmodulin-binding transcription activator 3-like |

| | | | | | |
|---|---|---|---|---|---|
| INS | chr07 | 26204256 | 26204257 | Vitvi07g01991 | putative MRP-like ABC transporter gene |
| DUP | chr09 | 923618 | 924383 | Vitvi09g01504 | uncharacterized |
| DUP | chr09 | 937904 | 940108 | Vitvi09g01505 | uncharacterized |
| DUP | chr09 | 11045993 | 11103730 | | |
| DUP | chr09 | 11052848 | 11053348 | Vitvi09g00849 | putative LRR receptor-like serine/threonine-protein kinase |
| DUP | chr09 | 11055831 | 11055989 | Vitvi09g00850 | putative LRR receptor-like serine/threonine-protein kinase |
| DUP | chr09 | 11056339 | 11056730 | Vitvi09g00851 | putative LRR receptor-like serine/threonine-protein kinase |
| DUP | chr09 | 11063100 | 11063710 | Vitvi09g00852 | putative LRR receptor-like serine/threonine-protein kinase |
| DUP | chr09 | 11064560 | 11064957 | Vitvi09g00853 | 7-dehydrocholesterol reductase |
| DUP | chr09 | 11075851 | 11076220 | Vitvi09g00854 | uncharacterized |
| DUP | chr09 | 11086261 | 11086779 | Vitvi09g00855 | cytochrome P450 CYP82D47-like |
| DUP | chr09 | 11089336 | 11092950 | Vitvi09g00856 | uncharacterized |
| DUP | chr09 | 11098014 | 11101644 | Vitvi09g00857 | cytochrome P450 CYP82D47-like |
| INS | chr10 | 20116908 | 20116909 | Vitvi10g01405 | uncharacterized |
| DEL | chr11 | 10498979 | 10499046 | | |
| DEL | chr11 | 11539526 | 11541421 | Vitvi09g00857 | histone-lysine N-methyltransferase ATXR3 |
| DEL | chr13 | 7312866 | 7312926 | | |
| DEL | chr14 | 16901078 | 16901221 | | |
| DEL | chr14 | 21038573 | 21038735 | | |
| DEL | chr15 | 6543232 | 6554147 | | |
| TRA | chr15 | 9713257 | 9713410 | | |
| DEL | chr16 | 19194133 | 19194199 | Vitvi16g01907 | short-chain dehydrogenase TIC 32 |
| DEL | chr16 | 19386663 | 19386914 | | |
| DEL | chr16 | 19736456 | 19736539 | | |
| DEL | chr17 | 13522597 | 13522672 | | |

| | | | | | |
|---|---|---|---|---|---|
| DEL | chr18 | 6349792 | 6350071 | | |
| DEL | chr18 | 17902281 | 17903662 | | |
| DEL | chr18 | 17902791 | 17903662 | Vitvi18g01415 | CTD nuclear envelope phosphatase 1 homolog |
| DEL | chr19 | 3198723 | 3198867 | | |
| DEL | chr19 | 4494970 | 4495022 | | |
| DEL | chr19 | 22299859 | 22299923 | | |

**Supplemental text 1**

FALCON-Unzip parameters for the Char04 genome.

```
[General]
input_fofn = 0-rawreads/input_preads.fofn
input_type = preads
length_cutoff = 3000
falcon_sense_skip_contained = TRUE
falcon_sense_option = --output_multi --min_idt 0.70 --min_cov 4 --max_n_read 400
length_cutoff_pr = 9000
pa_DBsplit_option = -x500
pa_HPCdaligner_option = -v -dal128 -t30 -e0.7 -M60 -l1000 -k16 -h64 -w7 -s1000 -T16
ovlp_DBsplit_option = -x500
ovlp_HPCdaligner_option = -v -B128 -M60 -t60 -k20 -h256 -e.9 -l1000 -s100 -T16
overlap_filtering_setting = --max_diff 100 --max_cov 400 --min_cov 3
```

**Supplemental text 2**

SRA Datasets of RNAseq experiments

- SRR1573044
- SRR1573043
- SRR1573042
- SRR1573041
- SRR1748359
- SRR1748358
- SRR1748357
- SRR1748356
- SRR1748355
- SRR1748354
- SRR1748353
- SRR1748352
- SRR1748351
- SRR1748350
- SRR1748349
- SRR1748348
- SRR1573044
- SRR1573043
- SRR1573042
- SRR1573041
- SRR1708660
- SRR1708662
- SRR1708663
- SRR1708899
- SRR1708900
- SRR1708901

ESTs

- Vitis ESTs (NCBI)
- Vitis vinifera cDNA cDNA Full Length (NCBI)

De novo assembled transcriptomes

- Venturini et al. - cv. Corvina - pool of 45 tissues

Proteins

- swissProt viridiplantae
- Vitis (NCBI)

Iso-Seq

- Cv. Cabernet Sauvignon (SRP132320)

**Supplemental text 3**

EVM weights

| | | |
|---|---|---|
| ABINITIO_PREDICTION | AUGUSTUS | 5 |
| ABINITIO_PREDICTION | GeneMark.hmm3 | 5 |
| ABINITIO_PREDICTION | augustus_BUSCOv3 | 6 |
| ABINITIO_PREDICTION | snap | 3 |
| PROTEIN | protein2genome | 1 |
| TRANSCRIPT | est2genome | 10 |
| TRANSCRIPT | PASA_assemblies | 20 |
| OTHER_PREDICTION | PASA_transdecoder | 20 |

**Supplemental text 4**

PASA parameters

```
#spliced multiple mappings allowed
run_spliced_aligners.pl:-N=5


#script validate_alignments_in_db.dbi
validate_alignments_in_db.dbi:--MIN_PERCENT_ALIGNED=90
validate_alignments_in_db.dbi:--MIN_AVG_PER_ID=95
validate_alignments_in_db.dbi:--MAX_INTRON_LENGTH=10000

#script subcluster_builder.dbi
subcluster_builder.dbi:-m=50

#annotation comparison
cDNA_annotation_comparer.dbi:--MIN_PERCENT_OVERLAP=75
cDNA_annotation_comparer.dbi:--MIN_PERCENT_PROT_CODING=40
cDNA_annotation_comparer.dbi:--MIN_PERID_PROT_COMPARE=70
cDNA_annotation_comparer.dbi:--MIN_PERCENT_LENGTH_FL_COMPARE=50
cDNA_annotation_comparer.dbi:--MIN_PERCENT_LENGTH_NONFL_COMPARE=50
cDNA_annotation_comparer.dbi:--MIN_PERCENT_ALIGN_LENGTH=50
cDNA_annotation_comparer.dbi:--MIN_PERCENT_OVERLAP_GENE_REPLACE=90
cDNA_annotation_comparer.dbi:--MAX_UTR_EXON
```

**Supplemental text 5**

TIGRFAM (15.0) : TIGRFAMs are protein families based on Hidden Markov Models or HMMs

SFLD (3) : SFLDs are protein families based on Hidden Markov Models or HMMs

SignalP_GRAM_NEGATIVE (4.1) : SignalP (organism type gram-negative prokaryotes) predicts the presence and location of signal peptide cleavage sites in amino acid sequences for gram-negative prokaryotes.

SUPERFAMILY (1.75) : SUPERFAMILY is a database of structural and functional annotation for all proteins and genomes.

PANTHER (12.0) : The PANTHER (Protein ANalysis THrough Evolutionary Relationships) Classification System is a unique resource that classifies genes by their functions, using published scientific experimental evidence and evolutionary relationships to predict function even in the absence of direct experimental evidence.

Gene3D (4.2.0) : Structural assignment for whole genes and genomes using the CATH domain structure database

Hamap (2017_10) : High-quality Automated and Manual Annotation of Microbial Proteomes

Coils (2.2.1) : Prediction of Coiled Coil Regions in Proteins

ProSiteProfiles (2017_09) : PROSITE consists of documentation entries describing protein domains, families and functional sites as well as associated patterns and profiles to identify them

SMART (7.1) : SMART allows the identification and analysis of domain architectures based on Hidden Markov Models or HMMs

CDD (3.16) : Prediction of CDD domains in Proteins

PRINTS (42.0) : A fingerprint is a group of conserved motifs used to characterise a protein family

ProSitePatterns (2017_09) : PROSITE consists of documentation entries describing protein domains, families and functional sites as well as associated patterns and profiles to identify them

Pfam (31.0) : A large collection of protein families, each represented by multiple sequence alignments and hidden Markov models (HMMs)

SignalP_EUK (4.1) : SignalP (organism type eukaryotes) predicts the presence and location of signal peptide cleavage sites in amino acid sequences for eukaryotes.

ProDom (2006.1) : ProDom is a comprehensive set of protein domain families automatically generated from the UniProt Knowledge Database.

MobiDBLite (1.0) : Prediction of disordered domains Regions in Proteins

PIRSF (3.02) : The PIRSF concept is being used as a guiding principle to provide comprehensive and non-overlapping clustering of UniProtKB sequences into a hierarchical order to reflect their evolutionary relationships.

SignalP_GRAM_POSITIVE (4.1) : SignalP (organism type gram-positive prokaryotes) predicts the presence and location of signal peptide cleavage sites in amino acid sequences for gram-positive prokaryotes.

TMHMM (2.0c) : Prediction of transmembrane helices in protein
